# Comparative genomics of *Xylella fastidiosa* suggests determinants of host-specificity and expands its mobile genetic elements repertoire

**DOI:** 10.1101/2021.10.17.464729

**Authors:** Guillermo Uceda-Campos, Oseias R. Feitosa-Junior, Caio R. N. Santiago, Paulo M. Pierry, Paulo A. Zaini, Wesley O. de Santana, Joaquim Martins-Junior, Deibs Barbosa, Luciano A. Digiampietri, João C. Setubal, Aline M. da Silva

## Abstract

The Gram-negative bacterium *Xylella fastidiosa* colonizes plant xylem vessels and is obligately vectored by xylem sap-feeding hemipteran insects. *X. fastidiosa* causes diseases in many plant species but in a variety of its plant hosts this bacterium behaves as a commensal endophyte. Originally confined to the Americas, infecting mainly grapevine, citrus and coffee plants, *X. fastidiosa* has spread to several plant species in Europe, causing devastating crop diseases. Although many pathogenicity and virulence factors have been identified in *X. fastidiosa* which enable the bacterium to successfully establish in the xylem tissue, the mechanisms by which distinct *X. fastidiosa* strains colonize and cause disease in specific plant hosts have not been fully elucidated. Here we present comparative analyses of 94 publicly available whole-genome sequences of *X. fastidiosa* strains with the goal of providing insights into plant host specificity determinants for this phytopathogen as well as of expanding the knowledge of its mobile genetic elements (MGE) content, mainly prophages. Our results revealed a pangenome of 4,549 protein coding sequences (CDSs) which is still open. The core- and accessory genomes comprise 954 and 2,219 CDSs, respectively. Phylogenetic tree construction using all core genome CDSs grouped the strains in three major clades of subspecies *fastidiosa, multiplex* and *pauca*, with subclades related to the strains’ sequence type (ST) obtained from multi-locus sequence typing (MLST). The geographic region where the strains were collected showed stronger association with the clades of *X. fastidiosa* strains rather than the plant species from which they were isolated. Among the CDS related to virulence and pathogenicity found in the core genome, those related to lipopolysaccharide (LPS) synthesis and trimeric autotransporter adhesins (TAA) are somewhat related with the plant host of a given strain according to phylogenetic inference. The *X. fastidiosa* accessory genome is represented by an abundant and heterogeneous mobilome, which includes a diversity of prophage regions. In summary, the genome comparisons reported here will enable a better understanding of the diversity of phylogenetically close genomes and warrant further investigation of LPS and TAAs as potential *X. fastidiosa* host-specificity determinants.

**Impact statement:** The bacterium *Xylella fastidiosa* is a pathogen that infects many plant species and has caused devastating diseases in grapevine, citrus, coffee, and olive plants. This phytopathogen *X. fastidiosa* is original from the Americas and has emerged in Europe where it is causing severe economic losses for olive producers, mainly in Italy. Although many pathogenicity and virulence factors have been identified in *X. fastidiosa*, which enable this bacterium to successfully establish in the xylem vessels network, the mechanisms by which distinct *X. fastidiosa* strains colonize and cause disease in the different plant host species have not been fully elucidated. The comparative analyses of 94 whole-genome sequences from *X. fastidiosa* strains from diverse hosts and geographic regions provide insights into host specificity determinants for this phytopathogen as well as expand the knowledge of its mobile genetic elements (MGE) content, mainly prophages. Our results contribute for a better understanding of the diversity of phylogenetically close genomes and warrant further experimental investigation of lipopolysaccharide and trimeric autotransporter adhesins as potential host-specificity determinants for *X. fastidiosa*.

**Data summary:** All genomic sequences were accessed from publicly available GenBank RefSeq database at NCBI (National Center for Biotechnology Information). A full listing of NCBI accession numbers for *X. fastidiosa* strains described in this paper is available in Table S1 (available in the online version of this article).

## Introduction

*Xylella fastidiosa* is a Gram-negative bacterium in the Xanthomonadaceae family that colonizes the xylem vessels of its plant hosts and is exclusively vectored by xylem sap-feeding hemipteran insects [1, 2]. This bacterium causes several crop diseases, such as Pierce’s disease (PD) of grapevine [3], citrus variegated chlorosis (CVC) [4], coffee leaf scorch (CLS) [5], plum leaf scald (PLS) [6], and olive quick decline syndrome (OQDS) [7]. While *X. fastidiosa* has also been associated with diseases in many other plant species, the bacterium behaves as a commensal endophyte in a variety of its plant hosts [8, 9].

A range of pathogenicity and virulence factors has been identified in *X. fastidiosa* that potentially enable the bacterium to overcome host defenses and successfully establish in the xylem tissue [1, 8, 10]. *X. fastidiosa* cells form biofilm-like structures that are crucial for successful acquisition and transmission by the insect vectors as well as for plant host colonization and pathogenesis [1, 11]. Progression of the disease symptoms are associated to *X. fastidiosa* systemic spread through the xylem vessel network which requires dispersal of bacterial cells from the biofilms [12–15] as well as twitching motility [16] and degradation of pit membranes by bacterial cell wall–degrading enzymes (CWDEs) [17, 18]. Moreover, symptoms severity is exacerbated by host-derived xylem occlusions (i.e., tyloses) elicited by *X. fastidiosa* colonization of grapevine [19]. Indeed, the symptoms caused by *X. fastidiosa* infection are suggestive of hydric stress and vary in intensity depending on pathogen genotype, plant host species/genotype, plant age, cultivation practices, and environmental conditions [10, 20].

Originally confined to the Americas, infecting mainly grapevine, citrus and coffee plants, *X. fastidiosa* has spread to various plants species in a number of European countries, possibly through the importation of infected plant material [8, 21, 22]. Currently, most of *X. fastidiosa* strains are categorized in three major subspecies *fastidiosa*, *pauca* and *multiplex* which are presumed to have originated in Central America (subsp. *fastidiosa*), South America (subsp. *pauca*) and North America (subsp. *multiplex*) [8, 9, 23]. Another two subspecies (subspp. *sandyi* and *morus*) native from North America have also been proposed [24, 25]. Furthermore, *X. fastidiosa* strains can be classified into sequence types (STs) based on a multilocus sequence typing (MLST) scheme with seven housekeeping genes [26, 27].

There is a loose association of *X. fastidiosa* subspecies or STs with host specificity, yet some strains can infect multiple hosts [10, 28]. Indeed, intersubspecific homologous recombination is known to drive *X. fastidiosa* adaptation to novel hosts [24, 29, 30]. However, the mechanisms by which the distinct *X. fastidiosa* strains successfully colonize specific plant hosts remain unclear. *X. fastidiosa* lacks the Type III secretion system (T3SS) [31], a membrane-embedded nanomachine typical of Gram-negative pathogens, which delivers effector proteins directly into host cells triggering or suppressing defense mechanisms, respectively in resistant or susceptible plants [32]. Instead, *X. fastidiosa* type II secretion system (T2SS) seems to be a relevant delivering source of its virulence proteins [10, 15, 33, 34]. It has been suggested that compatibility between xylem pit membrane carbohydrate composition and *X. fastidiosa* T2SS-secreted cell wall degrading enzymes is necessary for disease progression [35]. Moreover, since *X. fastidiosa* lipopolysaccharide (LPS) long chain O-antigen effectively delays plant innate immune recognition in grapevine, the heterogeneity of O-antigen composition may be among the mechanisms underlying *X. fastidiosa* host range [36].

Comparative genomics studies of *X. fastidiosa* strains isolated from different plant hosts and from diverse geographical regions identified shared and exclusive genes among these strains, chromosome rearrangements, indels, single nucleotide polymorphisms (SNPs) as well as differences in their mobile genetic elements (MGE) repertoire, such as plasmids, genomic islands and prophages [22, 29, 30, 37–48]. While some studies suggest that strains belonging to a phylogenetic group have similar pathogenicity mechanisms and strong selection, possibly driven by host adaptation, and, therefore, can be separated in subspecies [45, 46], other studies identified differences in each phylogenetic clade, such as enriched molecular functions [43] and distinct rates and events of recombination [22, 29, 30, 47].

The availability of new whole genome sequences of *X. fastidiosa* strains from diverse plant hosts and distinct geographical regions fosters up-to-date comparisons to be made. Here we present a comparative analysis of 94 *X. fastidiosa* genomes with the goal of providing insights into host specificity determinants for this phytopathogen as well as expanding the knowledge of its MGE content.

## Methods

### Data collection, curation, and annotation

A collection of 132 *X. fastidiosa* genome assemblies were downloaded from National Center for Biotechnology Information (NCBI) RefSeq database [49] (https://www.ncbi.nlm.nih.gov/genome/genomes/173) accessed in 2021-07-19 (Table S1). This initial collection was curated following the workflow depicted in Fig. S1 to remove genomes of laboratory variants, redundancies, and assemblies with contamination ≥5%, or with ≥1% of ambiguous bases, or with less than 20 tRNA genes or missing any of the 3 rRNA genes. Contamination and completeness of genome assemblies were evaluated using CheckM software [50]. Ambiguous bases in the assemblies were evaluated using QUAST tool [51]. Genomes that were not selected in the first curation round but represented a non-redundant strain, host or geographical region and had an associated publication were retrieved and included in the final curated collection, making a total of 94 genome assemblies (Table S1; Table 1). This final collection was annotated using Prokaryotic Genome Annotation Pipeline (PGAP) [52] standalone software package (https://github.com/ncbi/pgap), release 2021-07-01.build5508.

**Table 1.** Final curated collection of *Xylella fastidiosa* genome assemblies

| Strain name | Assembly accession | Genome assembly status | Plasmid number | Country of isolation | Host of origin | Sequence Type |
| --- | --- | --- | --- | --- | --- | --- |
| 32 | GCA_000506405 | Contig | 0 | Brazil | *Coffea sp* | 16 |
| 3124 | GCA_001456195 | Complete | 0 | Brazil | *Coffea arabica* | 16 |
| 11399 | GCA_001684415 | Contig | 2 | Brazil | *Citrus sinensis* | 11 |
| 6c | GCA_000506905 | Contig | 1 | Brazil | *Coffea sp* | 14 |
| 9a5c | GCA_000006725 | Complete | 2 | Brazil | *Citrus sinensis* | 13 |
| AlmaEM3 | GCA_018069645 | Complete | 0 | USA | *Vaccinium sp* | 42 |
| Ann-1 | GCA_000698805 | Complete | 1 | USA | *Nerium sp* | 5 |
| ATCC-35871 | GCA_000428665 | Scaffold | 0 | USA | *Prunus sp* | 41 |
| ATCC-35879 | GCA_011801475 | Complete | 1 | USA | *Vitis sp* | 2 |
| B111 | GCA_013283685 | Scaffold | 2 | Brazil | *Citrus sinensis* | 11 |
| Bakersfield-1 | GCA_009664125 | Complete | 1 | USA | *Vitis vinifera* | 1 |
| Bakersfield-11 | GCA_015476015 | Complete | 1 | USA | *Vitis vinifera* | 1 |
| Bakersfield-13 | GCA_015475995 | Complete | 1 | USA | *Vitis vinifera* | 1 |
| Bakersfield-14 | GCA_015475975 | Complete | 1 | USA | *Vitis vinifera* | 1 |
| Bakersfield-8 | GCA_015476035 | Complete | 1 | USA | *Vitis vinifera* | 1 |
| BB01 | GCA_001886315 | Scaffold | 0 | USA | *Vaccinium corymbosum* | 42 |
| BB08-1 | GCA_018069665 | Complete | 0 | USA | *Vaccinium sp* | 43 |
| CFBP7970 | GCA_004016315 | Contig | 1 | USA | *Vitis sp* | 2 |
| CFBP8071 | GCA_004016295 | Contig | 1 | USA | *Prunus dulcis* | 1 |
| CFBP8072 | GCA_001469345 | Scaffold | 0 | Ecuador | *Coffea arabica* | 74 |
| CFBP8073 | GCA_001469395 | Scaffold | 0 | Mexico | *Coffea canephora* | 75 |
| CFBP8078 | GCA_004016365 | Contig | 0 | USA | *Vinca sp* | 51 |
| CFBP8082 | GCA_004016375 | Contig | 1 | USA | *Ambrosia artemisiifolia* | 2 |
| CFBP8351 | GCA_004016405 | Contig | 1 | USA | *Vitis vinifera* | 1 |
| CFBP8356 | GCA_004016415 | Scaffold | 0 | Costa-Rica | *Coffea arabica* | 76 |
| CFBP8416 | GCA_001971475 | Contig | 0 | France | *Polygala myrtifolia* | 7 |
| CFBP8417 | GCA_001971505 | Contig | 0 | France | *Spartium junceum* | 6 |
| CO33 | GCA_001417925 | Contig | 0 | Costa-Rica | *Coffea sp* | 72 |
| CoDiRO | GCA_000811965 | Contig | 1 | Italy | *Olea sp* | 53 |
| COF0324 | GCA_001549815 | Contig | 4 | Brazil | *Coffea sp* | 14 |
| COF0407 | GCA_001549825 | Contig | 4 | Costa-Rica | *Coffea sp* | 53 |
| CVC0251 | GCA_001549765 | Contig | 4 | Brazil | *Citrus sinensis* | 11 |
| CVC0256 | GCA_001549745 | Contig | 4 | Brazil | *Citrus sinensis* | 11 |
| De-Donno | GCA_002117875 | Complete | 1 | Italy | *Olea europaea* | 53 |
| Dixon | GCA_000166835 | Scaffold | 1 | USA | *Prunus sp* | 6 |
| DSM-10026 | GCA_900129695 | Scaffold | 0 | USA | *Vitis vinifera* | 2 |
| EB92-1 | GCA_000219235 | Contig | 1 | USA | *Sambucus canadensis* | 1 |
| ESVL | GCA_004023385 | Contig | 2 | Spain | *Prunus dulcis* | 6 |
| Fb7 | GCA_001456335 | Complete | 1 | Argentina | *Citrus sp* | 69 |
| Fillmore | GCA_012974105 | Complete | 0 | USA | *Olea europaea* | 81 |
| GB514 | GCA_000148405 | Complete | 1 | USA | *Vitis sp* | 1 |
| Griffin-1 | GCA_000466025 | Contig | 0 | USA | *Quercus rubra* | 7 |
| GV156 | GCA_009910885 | Contig | 0 | Taiwan | *Vitis vinifera* | 2 |
| GV230 | GCA_014249995 | Complete | 0 | Taiwan | *Vitis labrusca* | 2 |
| Hib4 | GCA_001456315 | Complete | 1 | Brazil | *Hibiscus sp* | 70 |
| IAS-AXF212H7 | GCA_009669445 | Contig | 1 | Spain | *Prunus dulcis* | 6 |
| IAS-AXF-235T10 | GCA_009669465 | Contig | 1 | Spain | *Prunus dulcis* | 6 |
| IVIA5235 | GCA_003515915 | Complete | 1 | Spain | *Prunus avium* | 1 |
| IVIA5901 | GCA_004023395 | Complete | 0 | Spain | *Prunus dulcis* | 6 |
| IVIA6586-2 | GCA_009669335 | Contig | 2 | Spain | *Helicrysum italicum* | 6 |
| IVIA6731 | GCA_009669375 | Contig | 2 | Spain | *Helicrysum italicum* | 6 |
| J1a12 | GCA_001456235 | Complete | 2 | Brazil | *Citrus sp* | 11 |
| LM10 | GCA_012974145 | Complete | 0 | USA | *Olea europaea* | 7 |
| M12 | GCA_000019325 | Complete | 0 | USA | *Prunus sp* | 7 |
| M23 | GCA_000019765 | Complete | 1 | USA | *Prunus sp* | 1 |
| Ma151 | GCA_018449095 | Contig | 0 | Italy | *Rhamnus alaternus* | 87 |
| MUL0034 | GCA_000698825 | Complete | 1 | USA | *Morus alba* | 30 |
| Mul-MD | GCA_000567985 | Contig | 0 | USA | *Morus alba* | 29 |
| NOB1 | GCA_012952075 | Scaffold | 0 | USA | *Vitis rotundifolia* | 2 |
| OK3 | GCA_012952085 | Scaffold | 0 | USA | *Vitis vinifera* | 2 |
| OLS0478 | GCA_001549755 | Contig | 2 | Costa-Rica | *Nerium sp* | 53 |
| OLS0479 | GCA_001549735 | Contig | 4 | Costa-Rica | *Nerium sp* | 53 |
| PD7202 | GCA_006370235 | Contig | 0 | Netherlands | *Plant tissue* | undetermined |
| PD7211 | GCA_006370175 | Contig | 0 | Netherlands | *Plant tissue* | 73 |
| Pr8x | GCA_001456295 | Complete | 1 | Brazil | *Prunus sp* | 14 |
| RAAR14-plum327 | GCA_009695495 | Contig | 0 | Brazil | *Prunus domestica* | 26 |
| RAAR6-Butte | GCA_009695485 | Contig | 0 | USA | *Prunus dulcis* | 7 |
| Red-Oak-2 | GCA_015475935 | Complete | 0 | USA | *Quercus rubra* | 7 |
| RH1 | GCA_012974125 | Complete | 0 | USA | *Olea europaea* | 7 |
| Riv5 | GCA_015475955 | Complete | 1 | USA | *Prunus cerasifera* | 34 |
| Salento-1 | GCA_002954185 | Complete | 1 | Italy | *Olea europaea* | 53 |
| Salento-2 | GCA_002954205 | Complete | 1 | Italy | *Olea europaea* | 53 |
| Stag-s-Leap | GCA_001572105 | Contig | 1 | USA | *Vitis sp* | 1 |
| sycamore-Sy-VA | GCA_000732705 | Contig | 0 | USA | *Platanus occidentalis* | 8 |
| Temecula1 | GCA_000007245 | Complete | 1 | USA | *Vitis sp* | 1 |
| Temecula1Star | GCA_006370185 | Contig | 0 | USA | *Vitis sp* | 1 |
| TemeculaL | GCA_006370155 | Scaffold | 0 | USA | *Vitis sp* | 1 |
| TOS14 | GCA_007713995 | Contig | 0 | Italy | *Spartium junceum* | 87 |
| TOS4 | GCA_007713905 | Contig | 0 | Italy | *Prunus dulcis* | 87 |
| TOS5 | GCA_007713945 | Contig | 0 | Italy | *Polygala myrtifolia* | 87 |
| TPD3 | GCA_007845655 | Contig | 0 | Taiwan | *Vitis vinifera* | 2 |
| TPD4 | GCA_007845705 | Contig | 0 | Taiwan | *Vitis vinifera* | 2 |
| U24D | GCA_001456275 | Complete | 1 | Brazil | *Citrus sinensis* | 13 |
| VB11 | GCA_012952095 | Scaffold | 1 | USA | *Vitis vinifera* | 2 |
| WM1-1 | GCA_006370215 | Contig | 0 | USA | *Vitis sp* | 2 |
| XF3348 | GCA_009669515 | Contig | 1 | Spain | *Prunus dulcis* | 81 |
| XRB | GCA_013283695 | Scaffold | 2 | Brazil | *Citrus sp* | 11 |
| XYL1732 | GCA_003973705 | Contig | 0 | Spain | *Vitis sp* | 1 |
| XYL1752 | GCA_009669505 | Contig | 1 | Spain | *Prunus dulcis* | 81 |
| XYL1968-18 | GCA_014856935 | Contig | 0 | Spain | *Olea europaea* | 81 |
| XYL1981 | GCA_009669455 | Contig | 0 | Spain | *Ficus carica* | 81 |
| XYL2055 | GCA_003973695 | Contig | 0 | Spain | *Vitis sp* | 1 |
| XYL2107-18 | GCA_014856795 | Scaffold | 0 | Spain | *Prunus dulcis* | 1 |
| XYL2153-18 | GCA_014856785 | Scaffold | 0 | Spain | *Vitis vinifera* | 1 |

### Genome comparisons

Comparative genomics analyses, pangenome, core genome and accessory genome reconstruction were performed using the Gene Tags Assessment by Comparative Genomics (GTACG) framework (https://github.com/caiorns/GTACG-backend). GTACG is based on an algorithm that uses clustering coefficient to find and maximize the number of orthologous groups in genomes from closely related strains [53]. The PGAP annotated genomes were uploaded in GTACG framework, and the protein coding sequences (CDSs) were compared using standalone BLASTP tool [54] with an e-value threshold of 1e-10. The clustering tool in GTACG framework was used to find a threshold that maximizes the cluster coefficient of each cluster. We found that a threshold of 45% of the alignment length was enough to produce concise homologous clusters. Metadata information of the *X. fastidiosa* strains (Table S1) such as plant host, country of isolation and sequence type (ST) were retrieved from NCBI RefSeq database, public databases for molecular typing and microbial genome diversity (PubMLST) [27] and from the literature, manually curated and analyzed together with the information provided by the GTACG framework.

### Phylogenetic analyses

Amino acid sequences of core genome orthologous CDSs were aligned with Clustal Omega v.1.2.1 [55] using default parameters. For maximum-likelihood (ML) phylogenies, the alignments were concatenated and computed using IQ-TREE v.1.5.4 [56] with a model predicted by ModelFinder and an ultrafast bootstrap of 1,000 replicates [57].

### Functional Annotation

Orthologous protein clusters encoded by the core, accessory and singleton genomes were compared to the Clusters of Orthologous Groups (COGs) [58] database using rpsblast+ (BLAST version 2.9.0) [54], with a cut-off e-value of 1e-6. COG categories were assigned to the best hits of rpsblast+ analysis.

### Mobile genetic elements prediction

Mobile Genetic Elements (MGE), such as prophages, genomic islands (GI) and insertion sequences (IS) were identified in the genome assemblies by a combination of prediction tools coupled with manual curation as previously described [59]. Prophage regions were predicted with Virsorter2 [60] and PHASTER [61]. Inovirus_detector software (https://github.com/simroux/Inovirus) was used for identification of prophages from the Inoviridae family (filamentous single-stranded DNA phages) [62]. GI consensus regions were defined using the results of SeqWord Sniffer [63] and GIPSy [64] software, which was used to assign one or more categories related to GI potential function. GI regions overlapping to prophages sequences were not considered. IS regions were predicted using the ISEScan [65] software. MGE regions predicted in each genome assembly were mapped in the genome for visual inspection and manual curation. Nucleotide sequences of prophages, GIs and ISs were compared to explore homology relationships using BLAST all-vs-all. The BLAST hits with an identity and coverage alignment higher than 40% and 75%, respectively, were filtered, analyzed and the resulting sequence similarity network (SSN) was visualized with Cytoscape 3.8 software [66]. Finally, the most frequent prophages and genomic islands were retrieved for the evaluation of their gene content. Taxonomic classification of selected prophages was performed with vContact2 [67] and with PhaGCN [68].

### Prospection of anti-MGE defense systems

CRISPR-Cas systems were searched with the software CRISPRFinder (http://crispr.i2bc.paris-saclay.fr/Server/) [69]. Hidden Markov Models (HMM) matrices were built to analyze known antiphage defense systems such as superinfection exclusion (SIE), Disarm, Brex, pAgos, Abortive Infection (Abi), Hachiman, Shedu, Septu, Lamassu, Druantia, Gabyja, Zorya and Wadjet [70]. To create HMM matrices, we recovered FASTA files with the amino acid sequences of each system from NCBI and IMG/M (Integrated Microbial Genomes & Microbiomes) databases [71] and created an alignment for a set of sequence of each system, which was then compared against the *X. fastidiosa* genomes. PICI elements (Phage-inducible chromosomal islands) were searched in *X. fastidiosa* genomes using an in-house Python pipeline that enables detection of the main PICI features [72]. Restriction-modification (R-M) systems were searched with BLASTP against the REBASE [73] database.

## RESULTS AND DISCUSSION

### General features of *X. fastidiosa* genomes

The main features of genome assemblies as well as plant host and country of isolation of 132 *X. fastidiosa* strains publicly available until 2021-07-19 in NCBI RefSeq database are summarized in Table S1. This collection was curated following the pipeline depicted in Figure S1 (to remove redundancies as well as genomes of laboratory variants) and 94 genome assemblies were selected for further analyses (Table S1; Table 1). These are high-quality draft genome sequences [74] given they present high completeness (>98%) and low contamination (<1.45%) according to CheckM [50] analysis. The average chromosome size of the selected 94 assemblies is 2,537,252 bp ± 90,235 bp with an average GC content of 51.88%± 0.36. Strains Hib4 (isolated from *Hibiscus* spp.) and Griffin-1 (isolated from *Quercus rubra*) have, respectively, the largest (2,813,286 bp) and smallest (2,387,314 bp) chromosome sizes. While 46 strains of the selected genome assemblies do not include plasmid related-contigs, the number of plasmids in the other strains is 1 (34 strains), 2 (9 strains), and 4 (5 strains), which include conjugative and mobilizable as well as non-mobilizable plasmids [42]. Chromosomes of the selected genomes have 2,291 ± 131 CDS and 110 ± 45 protein coding pseudogenes annotated by PGAP [52]. These results indicate a reasonable homogeneity in the genomes of distinct *X. fastidiosa* strains in relation to their chromosome sizes and GC content. In contrast, the plasmid content shows a greater diversity among strains consistent with previous observations [42].

### *X. fastidiosa* pangenome and core genome

The pangenome of *X. fastidiosa* (number of orthologous CDSs clusters present in the 94 genomes) was calculated using GTACG framework [53], considering chromosome and plasmids CDSs, since pangenomes are composites of the host chromosome together with MGEs [75]. The pangenome growth curve has not yet reached saturation (Fig.1a), indicating that the *X. fastidiosa* pangenome can be considered open and comprises 4,549 orthologous CDSs. The core genome curve (Fig. 1b) reveals that 954 CDSs belong to the core genome (conserved orthologous CDSs present in all 94 genomes). The pangenome frequency plot (Fig. 1c) shows the typical U-shape where 30.25% and 20.97% of pangenome p CDSs are detected, respectively, in a single genome (singleton genome) and in all genomes (core genome). Calculation of the soft-core genome (conserved orthologous CDSs present 95% of the selected genomes, i.e., 89 genomes) showed 1,567 CDSs (34.4% of the pangenome).

**Fig. 1.**
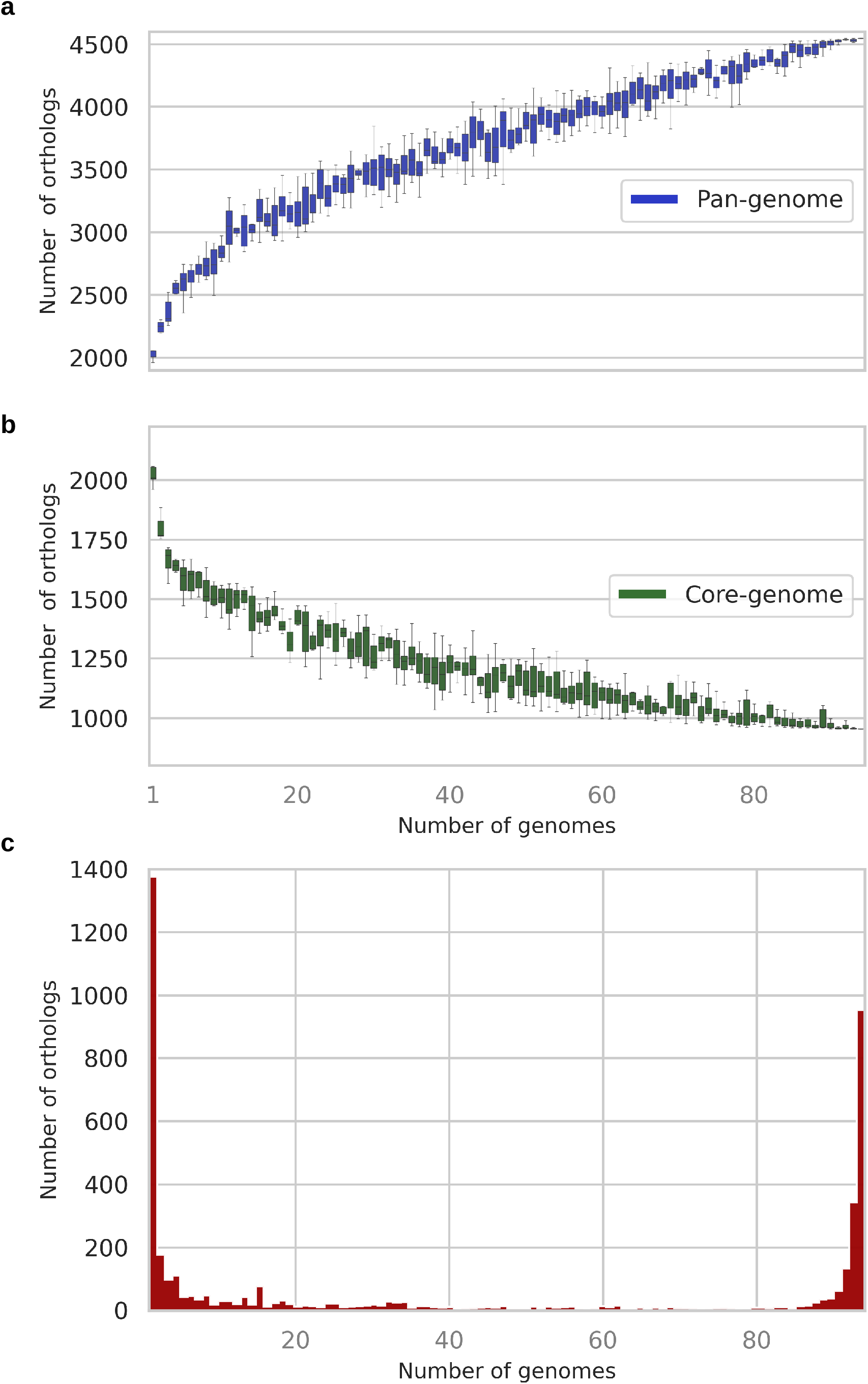
Pangenome and core-genome of 94 *Xylella fastidiosa* strains. Pangenome (a) and core genome (b) curves. Each boxplot represents the distribution of the number of orthologous CDSs clusters added (pangenome) or in common (core genome) with the addition of new genomes. Pangenome frequency plot (c). Number of orthologous CDSs detected in a single genome (left) and in all analyzed genomes (right).

The values for core genome as well as the pangenome frequency values we report here are somewhat different than previously reported [30, 43, 48] because we have used different algorithms for genome annotation and clustering of orthologous CDSs as well as a larger number of genomes.

### Genome-scale phylogeny

The core genome (954 CDSs) was used for a genome-scale phylogeny. The Maximum Likelihood (ML) tree (Fig. 2) grouped the 94 *X. fastidiosa* strains in three major clades defined by strains from the subspecies *fastidiosa, multiplex* and *pauca*. The strains from subspecies *morus* and *sandyi* grouped in subclades of the major *subsp. fastidiosa* clade. The overall topology of this core-genome based phylogeny tree agrees with a previously reported genome-wide phylogeny of 21 *X. fastidiosa* strains [45] and a *k-mers* based phylogeny of 72 *X. fastidiosa* strains [30].

**Fig. 2.**
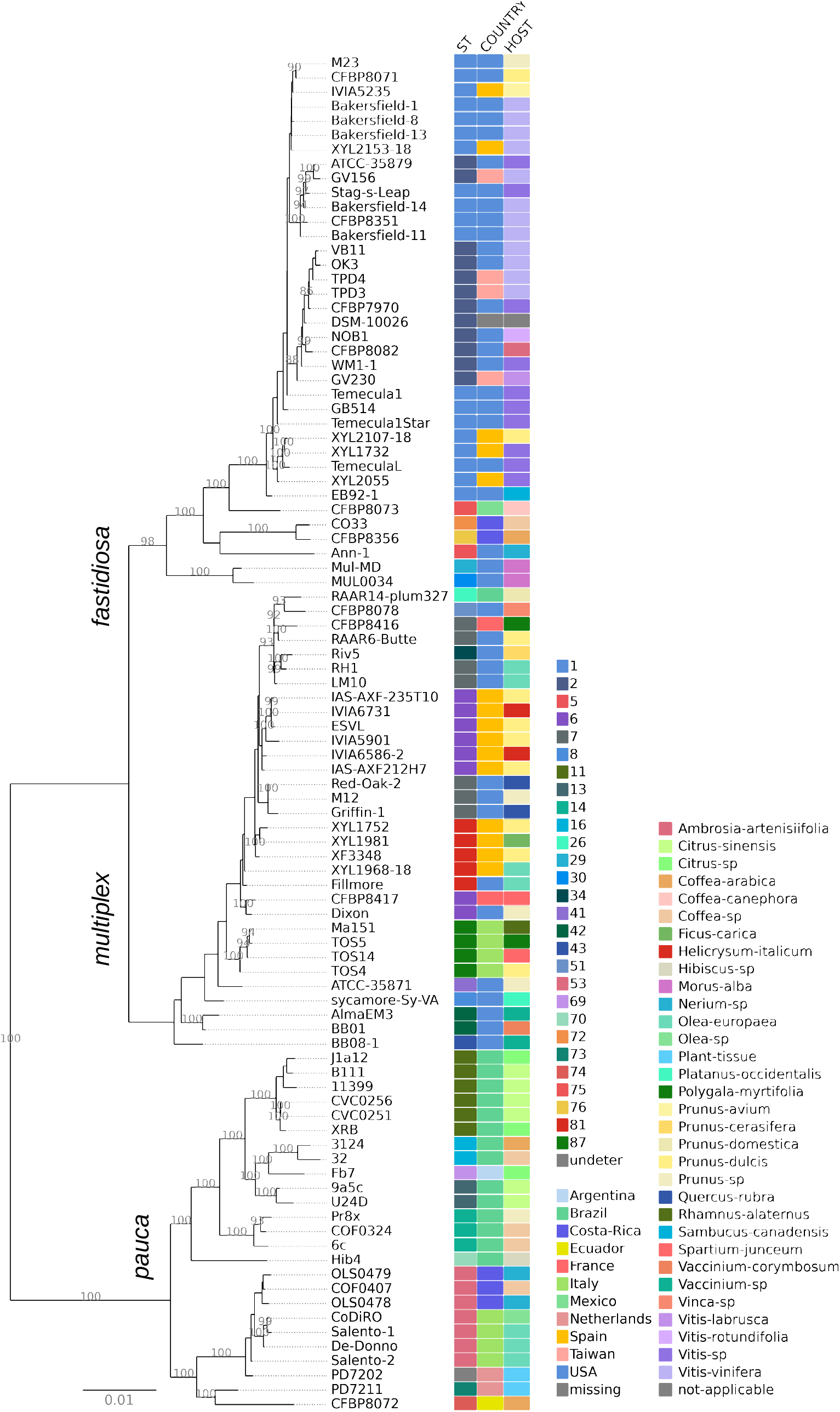
Core genome-scale phylogeny. Amino acid sequences of 954 CDSs of *X. fastidiosa* (94 strains) core genome were used for a Maximum Likelihood (ML) phylogenetic reconstruction. The three major clades grouped strains from subspecies *fastidiosa*, *multiplex* and *pauca*. Information of sequence type (ST), country of isolation and host of origin for each strain (Table 1) are represented by the squares followed by the indicated color legends.

Information of ST, country of isolation and host of origin for each strain (Table S1) were integrated to the genome-scale phylogeny (Fig. 2) as an attempt to highlight correlations, if any, among strain features and their phylogenetic relationship. We observe that most of the subclades are congruent with groups of the STs as well as country of isolation. For example, strains of ST1 belong to subclades of subsp. *fastidiosa* major clade and have been isolated in USA and Spain. Both ST6 and ST7 strains are in subclades of subsp. *multiplex* along with strains from USA, Spain and France. ST11, ST14 and ST53 were distributed among strains of subspecies *pauca*, which the first two STs are from strains isolated in Brazil while ST53 strains were isolated from Costa Rica and Italy. The strains from Italy were grouped with Costa Rica strains, corroborating the reported introduction of *X. fastidiosa* in Italy originating from Costa Rica [76]. Similarly, IVIA5235 (ST1; subsp. *fastidiosa*) isolated in Spain was possibly imported from North America as previously suggested [22]. In the case of STs represented by a single strain, most of them, such as ST5 (Ann-1), ST8 (sycamore-Sy-VA), ST43 (BB08-1), ST69 (Fb7), ST70 (Hib4), ST74 (CFBP8072) and ST76 (CFBP8356), are found in a branch by themselves.

The core-based phylogeny indicates a weak association between host of origin with the major clades in the genome-scale ML tree (Fig.2). Some strains isolated from *Coffea*, *Citrus*, *Olea*, *Vitis*, and *Morus* belong to monophyletic clades. It has been shown that citrus and coffee strains from subspecies *pauca* seem to be limited to their original hosts, despite crop proximity and the presence of insect vectors [77, 78]. On the other hand, the core-based phylogeny also indicates that some strains isolated from *Coffea*, *Prunus*, and *Nerium* are distributed into the three distinct major clades. There is evidence that some strains can infect multiple hosts [28, 79, 80] and that intersubspecific homologous recombination drives *X. fastidiosa* adaptation to novel hosts [24, 29, 30].

### Virulence factors as potential host specificity determinants

We found that the vast majority (90%; 63/70) of the CDSs listed in Table S2, which were identified or predicted to be virulence and pathogenicity factors for *X. fastidiosa* [10, 34, 36, 38, 81–84], belong either to the core or soft-core genomes. The lack of CDSs in some strains is mostly due to pseudogenization (data not shown). We highlight the polygalacturonase (PD1485 in Temecula1 strain) ortholog, previously reported to carry a frameshift mutation [38], which is confirmed as a pseudogene in strains from subspecies *pauca* isolated from citrus (strains 9a5c, U24D, Fb7, J1a12, B111, CVC0251, CVC0256, 11399 and XRB), coffee (strains 32 and 3124), and vinca (strain CFBP8078). All other strains from subspecies *pauca* such as Pr8x, 6c, Hib4, COF0324, CFBP8072, CODIRO and De-Donno harbor an intact polygalacturonase sequence, similarly to all other strains analyzed in this study from subsp. *multiplex* and *fastidiosa*. Polygalacturonase has been shown to be a critical virulence factor for *X. fastidiosa* pathogenesis in grapevine [18]; therefore, we hypothesize that another cell wall-degrading enzyme, such as a putative pectin-lyase [85], may perform that function in the strains that carry the frameshift mutation.

Each of the orthologous clusters of CDSs related to virulence/pathogenicity (Table S2) that belong to core or soft-core genomes was used for a separate phylogeny reconstruction. The resulting ML trees were inspected to verify evidence, if any, of association of clades and subclades with specific plant hosts. Among dozens of trees, we found that a few may reflect the kind of association we were looking for, such as the trees reconstructed with CDSs related to LPS biosynthesis and CDSs of the three trimeric autotransporter adhesins (TAA) (Fig.3)

**Fig. 3.**
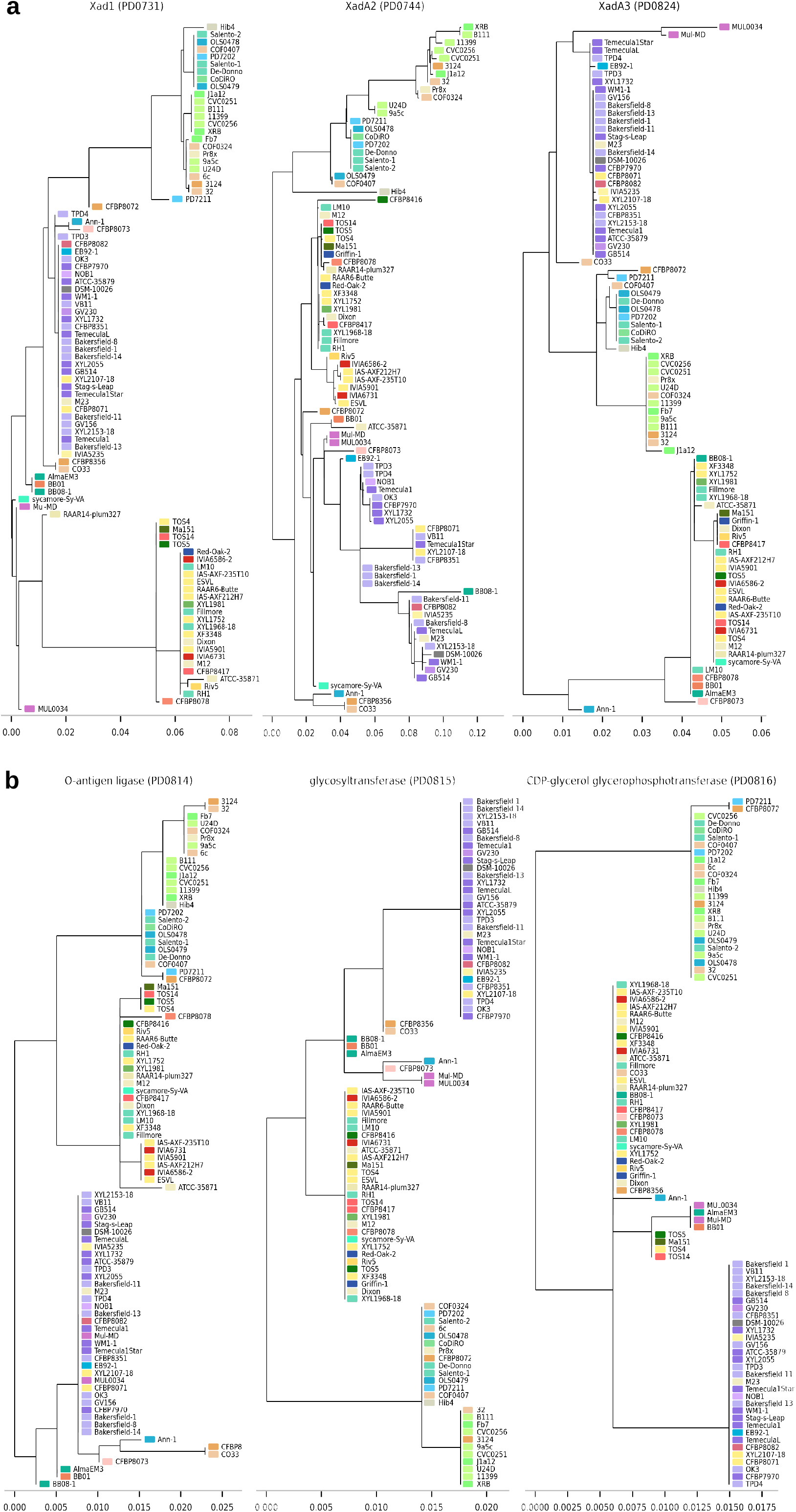
Phylogeny reconstruction of selected CDS. Maximum Likelihood (ML) phylogenetic reconstruction of CDS of three trimeric autotransporter adhesins (TAA) (a) and CDS related to LPS biosynthesis (b). Information of host of origin for each strain (Table 1) represented by the squares follow the indicated color legend shown in Fig. 2.

The ML trees (Fig. 3a) obtained with orthologous clusters of CDSs encoding the afimbrial adhesins *xadA1* (PD0731), *xadA2* (PD0744) and *xadA3* (PD0824) suggest that these genes, particularly *xadA1* and *xadA3*, are potential determinants of host specificity. These afimbrial adhesins mediate *X. fastidiosa* cell-cell aggregation and adhesion to surfaces during biofilm formation [11, 86]. The orthologs of PD0814 (O-antigen ligase family protein), PD0815 (Glycosyltransferase family 2 protein) and PD0816 (CDP-glycerol glycerophosphotransferase family protein), which are related to LPS biosynthesis, generated ML trees (Fig. 3b) that also suggest these genes, particularly the O-antigen ligase, as potential determinants of host specificity. It has been shown that O-antigen delays plant innate immune recognition in grapevine and as such the heterogeneity of O-antigen composition may be related to *X. fastidiosa* host range [36]. Overall, our results suggest that differences in the sequences of virulence-related genes may contribute to define *X. fastidiosa* host-specificity.

### Unraveling *X. fastidiosa* accessory genome and its mobile genetic elements

The distribution of core, singleton and accessory genomes of the 94 strains among COG functional categories is depicted in Fig. 4. As expected, the COG functional categories of highly conserved biological processes, such as “Translational, ribosomal structure, and biogenesis” (category J), and “Cell wall/membrane/envelope biogenesis” (category M) comprise a substantial fraction of the core genome in comparison to the accessory genome. In contrast, the accessory genome is enriched in category X (Mobilome: prophages, transposons), comprising ~15%. Other categories also enriched in the accessory genome are “Replication, recombination and repair” (category L) and “Defense mechanisms” (category V) which is suggestive of the ability of *X. fastidiosa* strains to cope with stress conditions in distinct environments.

**Fig. 4.**
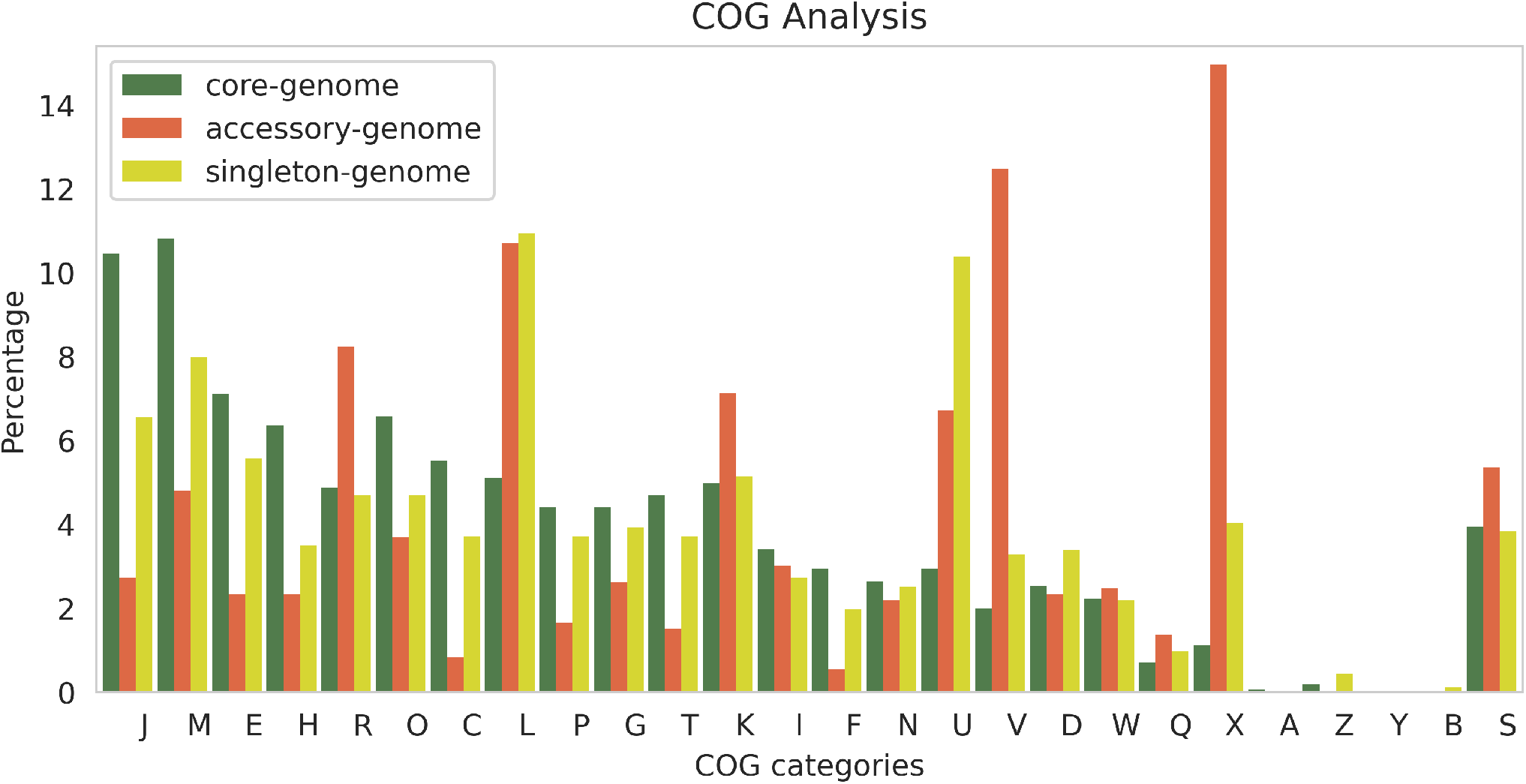
Distribution of core, singleton and accessory genomes of the 94 genomes strains among Clusters of Orthologous Groups (COG) functional categories. [J] Translation, ribosomal structure and biogenesis; [M] Cell wall/membrane/envelope biogenesis; [E] Amino acid transport and metabolism; [H] Coenzyme transport and metabolism; [R] General function prediction only; [O] Posttranslational modification, protein turnover, chaperones; [C] Energy production and conversion; [L] Replication, recombination and repair; [P] Inorganic ion transport and metabolism; [G] Carbohydrate transport and metabolism; [T] Signal transduction mechanisms; [K] Transcription; [I] Lipid transport and metabolism; [F] Nucleotide transport and metabolism; [N] Cell Motility; [U] Intracellular trafficking, secretion, and vesicular transport; [V] Defense mechanisms; [D] Cell cycle control, cell division, chromosome partitioning; [W] Extracellular structures; [Q] Secondary metabolites biosynthesis, transport and catabolism; [X] Mobilome: prophages, transposons; [A] RNA processing and modification; [Z] Cytoskeleton; [Y] Nuclear structure; [B] Chromatin structure and dynamics; [S] Function unknown.

The enrichment of the accessory genome in the mobilome-associated CDSs (COG category X) prompted us to explore the full set of MGEs (prophages, genomic islands, insertion sequences) in the genome assemblies of the 94 *X. fastidiosa* strains. Using a combination of prediction tools, sequences related to prophages, GIs, ISs, and plasmids were identified in the genome assemblies. We found that the content of MGEs varies considerably among the strains, ranging from ~5% to ~40% of the genome, with a mean value of 19.2% ± 8.3. Among the strains with the higher MGE content are RH1, J1a12, Fb7 and MUL0034 (Fig. 5).

**Fig. 5.**
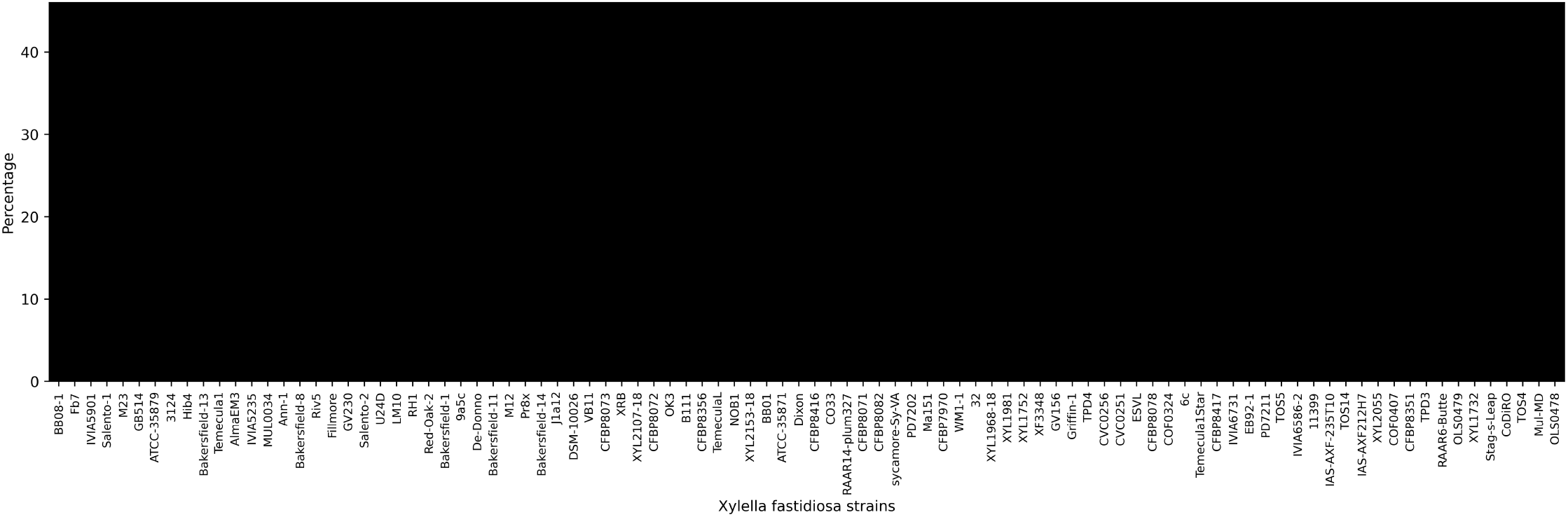
Percentage of mobile genetics elements distributed among the *X. fastidiosa* strains according to their genome assembly level: complete, scaffold and contig.

*X. fastidiosa* genome assemblies harbor 9 ± 2 prophage-related regions. Among the strains with complete genomes, IVIA5901, Hib4, MUL0034, and RH1 have the greatest number of prophage regions (10 regions) while the strains with the least prophage regions are Salento-1, Salento-2, De-Donno (5 regions). A previous study reported 6 and 8 prophage-like regions in complete genomes of 9a5c and Temecula1, respectively [87] and a comparison of 72 *X. fastidiosa* genomes revealed an average of 9.5, 9.3 and 8.5 prophage regions, respectively, for strains from subsp. *fastidiosa*, *multiplex* and *pauca* [30].

The MGEs identified in the genome assemblies of the 94 strains were then grouped in a sequence similarity network (SSN). Fig. 6 shows the clusters representing the predicted *X. fastidiosa* mobilome. While some sequences are conserved in various strains (clusters in Fig. 6) several are unique to a particular strain (shown in the bottom of Fig. 6). The sizes of these MGE sequences vary from ~4 kbp to 100 kbp for prophages and genomic islands, 100 bp to 4.8 kbp for insertion sequences, and 1 kbp to 64.3 kbp for plasmids (data not shown). Most of the MGEs clusters are from GIs with an average size of 23.7 kbp ± 11. A few GIs seem to be related to pathogenicity/virulence or to antibiotic resistance, such as cluster 1, cluster 10, and cluster 17, which harbor CDSs encoding efflux RND transporter and toxin-antitoxin systems. ISs appear distributed mainly in six clusters with tightly connected nodes (clusters 4, 9, 12, 13, 15, and 16) showing ISs commonly found among *X. fastidiosa* genomes. Several ISs of the clusters 4 and 8 are found within other MGEs such as prophages or genomic islands, while the other ISs were found in the chromosome. The ISs from clusters 4 and 8 belong to the IS200/IS605 family which is widely spread in Bacteria and Archaea [88]. Members of this family are unusual because they use obligatory single-strand DNA intermediates, which distinguishes them from classical IS [88].

**Fig. 6.**
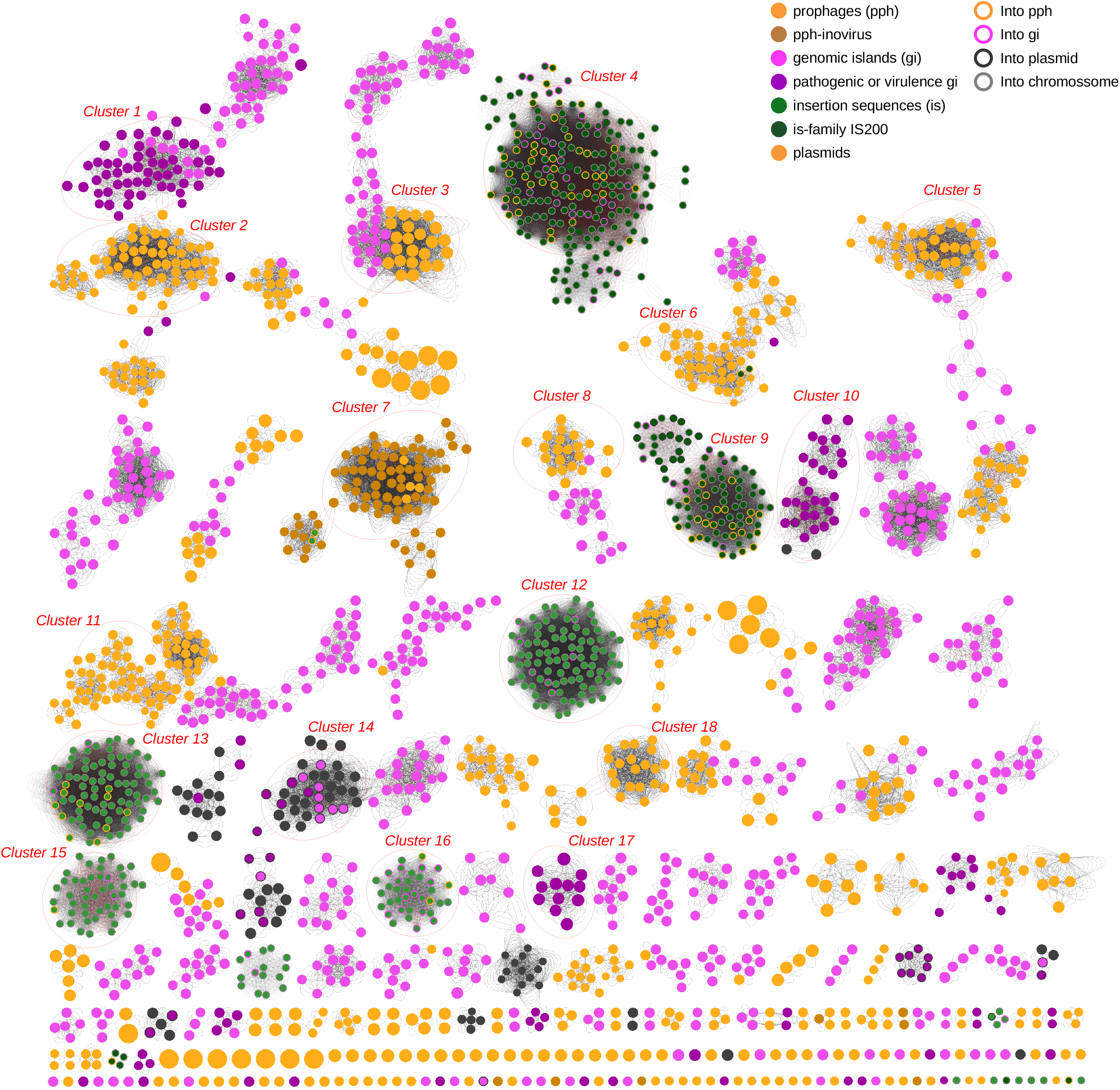
Sequence Similarity Network of the *X. fastidiosa* mobilome. Prophage, genomic islands, insertion sequences and plasmids as indicated by distinct colors. Nodes correspond to each of the distinct MGEs predicted in the 94 strains analyzed, the edges represent the similarity of nucleotide sequence with more than 75% of identity and 40% of the coverage, and evalue < 1e-6.

A closer examination of clusters 2, 8, 11, and 18 (Fig. 6) reveals that their prophage sequences carry lysozyme and holin proteins, commonly found in temperate and lytic bacteriophages. The sequences grouped in these 4 clusters belong from strains isolated from diverse countries such as Brazil, Mexico, Costa Rica, Italy, Spain and USA, and also in Taiwan in the case of clusters 2 and 11.

Cluster 7 groups prophages classified as inoviruses [62] and identified in 68 of the 94 genomes analyzed. Some inoviruses are present in two copies in a same strain such as Salento-1 and Salento-2 which could suggest superinfection events. It remains to be investigated whether multiple prophage carriage confers any fitness advantage to *X. fastidiosa*, as has been observed for *Pseudomonas aeruginosa*, where multiple prophage carriage seems to be beneficial during mixed bacterial infections [89]. Inoviruses play a relevant role in the structure in *P. aeruginosa* biofilm [90] and have been reported to encode *Zonula occludens* toxin (Zot) in several *Vibrio* species [91]. Zot protein seems to play a dual function as it is essential for inovirus morphogenesis and has also been reported to contribute for *Vibrio cholerae* pathogenesis [92, 93]. Zot-like CDSs are annotated in multiple inoviruses distributed among *X. fastidiosa* strains (data not shown). Zot proteins have been postulated as virulence factors for plant pathogens [94], including *X. fastidiosa* [41]. It is worth noting that EB92-1, a proposed *X. fastidiosa* biocontrol strain, lacks both Zot genes found in Temecula1 strain (PD0915 and PD0928) and as such Zot has been suggested as a potential *X. fastidiosa* virulence factor [95]. Moreover, a *X. fastidiosa* Zot protein was shown to elicit cell death-like responses in the apoplast of some *Nicotiana tabacum* cultivars [33]. Besides Zot, other prophage-encoded genes may play a role in the biology of *X. fastidiosa* as observed in other bacteria, where the so called “moron” loci have been related to virulence, stress resistance, phage resistance and host adaptation [96–98]. More studies are necessary to understand the contribution of “moron” loci, such as Zot genes, as well as events of prophage induction to *X. fastidiosa* biology. There is experimental evidence *X. fastidiosa* releases phage particles [99, 100] but the impact of prophage induction in host colonization is unknown.

### Immunity systems prospection in *X. fastidiosa* genomes

Since *X. fastidiosa* strains harbor numerous MGE, we made a screening of the well-known immunity systems in Gram-negative bacteria to evaluate *X. fastidiosa* strategy to deal with mobile genetic elements. Figure 7 shows the screening results for 46 *X. fastidiosa* genome assemblies. The SuperInfection Exclusion (SIE), Abortive infection, pAGOs, DISARM and BREX systems [101–105] are absent in all *X. fastidiosa* strains analyzed in this study. The same was observed for the recently reported systems HACHIMAN, SHEDU, SEPTU, LAMASSU and DRUANTIA [70]. Although we have found genes coding for proteins of the systems GABYJA and ZORYA [70] in all *X. fastidiosa* strains analyzed, none were inside an operon, and as such cannot be considered as true systems. The proteins gp41, gp42 and gp43 previously described as part of a SIE system operon in *X. fastidiosa* strain 53 [100] are found in several of the strains we have analyzed, although not as a complete operon and also not considered as true systems. We created HMM clusters and used them against all strains genomes searching for phage-inducible chromosomal islands (PICI) elements [72]. Although they are commonly found in Gram-negative bacteria, our analysis did not detect PICI elements in *X. fastidiosa* genomes.

**Fig. 7.**
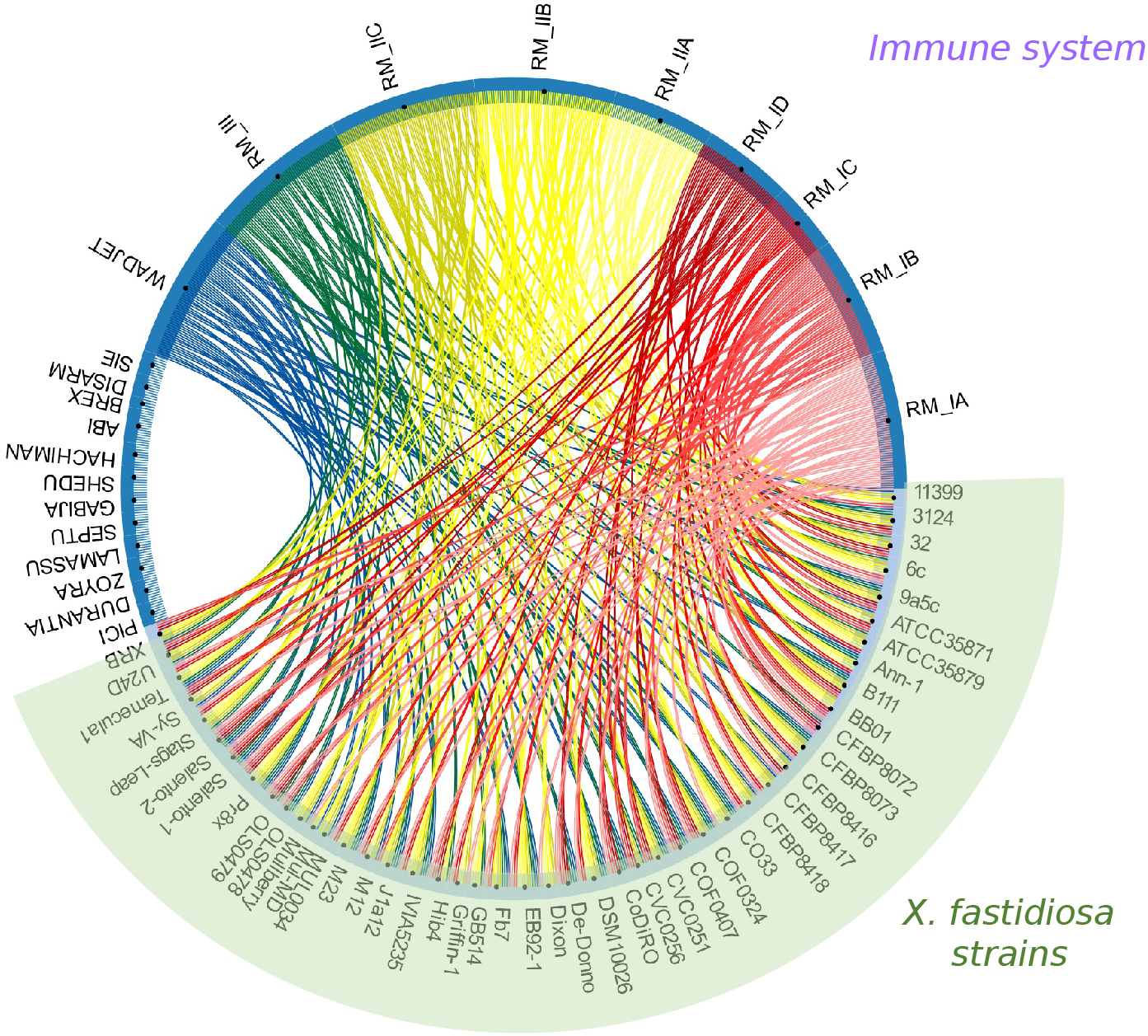
Circus plot showing the distribution of the different immunity systems in the genomes of *X. fastidiosa* strains.

A survey of Restriction-Modification systems (R-M system types I, II and III) [73, 106] in the 46 genome assemblies showed that all strains possess at least one of the three main R-M system types (Fig. 7) as previously reported for 9a5c and Temecula1 strains [107]. The type II is usually found in multiple operons per genome, while the type III is observed in a single operon per genome. We also searched for CRISPR sequences and genes encoding Cas proteins [69]. All potential CRISPR candidates found in the 94 genomes assemblies are not true CRISPRs according the CRISPRfinder tool [69]. Moreover, although genes encoding Cas-like proteins were found in some *X. fastidiosa* strains, they are not in the vicinity of any CRISPR candidates. Cas proteins are required for the functionality of the CRISPR/Cas immunity system [108]. Thus, similarly to major bacterial lineages [109], *X. fastidiosa* lacks a functional CRISPR-Cas viral defense system, which may contribute its permissiveness in prophage acquisition. Moreover, despite the fact that *X. fastidiosa* genomes encode R-M systems, a mechanism of immunity known to prevent both lytic and lysogenic infections in individual bacteria, it is reported to increase the number of prophage-acquiring individuals at the population level [110].

We also investigated the presence of the WADJET system reported to act against foreign plasmidial DNA [70]. This system was found in most of the *X. fastidiosa* strains we analyzed, except in ATCC35879, OLS0479, CVC0256 and 6c (Fig. 7). Strains OLS0479, CVC0256 do not have WADJET system, which may contribute to harboring 4 plasmids each.

### Final remarks

The comparative analyses of 94 publicly available whole-genome sequences of *X. fastidiosa* strains revealed an open pangenome with 4,549 protein coding sequences (CDS). A core genome-scale phylogeny grouped these *X. fastidiosa* strains in three major clades defined by strains from the subspecies *fastidiosa, multiplex* and *pauca* consistent with previous *k-mers* based phylogeny of 72 *X. fastidiosa* strains [30]. Most of the subclades are congruent with groups of the STs as well as country of origin. Moreover, the geographic region where the strains were collected showed stronger association with the clades of *X. fastidiosa* strains rather than the plant species from which they were isolated. The vast majority of the CDSs identified or predicted to be virulence and pathogenicity factors for *X. fastidiosa* belong either to the core or soft-core genomes. Among the CDS related to virulence and pathogenicity found in the core genome, those related to lipopolysaccharide (LPS) synthesis and trimeric autotransporter adhesins (TAA) are somewhat related with the plant host of a given strain according to phylogenetic inference, and as such may contribute to define *X. fastidiosa* host specificity. Finally, we found that the content of MGEs varies considerably among the strains, ranging from ~5% to ~40% of the genome assemblies and includes a diversity of sequences related to prophages, GI, IS and plasmids. It is worth noting the inoviruses sequences are found in all analyzed strains and that they encode a Zot protein which has been suggested to be a virulence factor for *X. fastidiosa*.

Overall, the comparative analyses of 94 whole-genome sequences from *X. fastidiosa* strains from diverse hosts and geographic regions provide insights into host specificity determinants for this phytopathogen as well as expand the knowledge of its mobile genetic elements (MGE) content. Our results contribute for a better understanding of the diversity of phylogenetically close genomes and warrant further experimental investigation of lipopolysaccharide and trimeric autotransporter adhesins as potential host-specificity determinants for *X. fastidiosa*.

## Authors Contributions

Conceptualization: A.M.D.S., G.U.C., O.R.F.J. Methodology: C.R.N.S., G.U.C., J.C.S., J.M.J., L.A.D., O.R.F.J. Computing resources: C.R.N.S., J.C.S., L.A.D. Data curation: C.R.N.S., G.U.C., O.R.F.J. Formal analysis: A.M.D.S., D.B., G.U.C., J.M.J., O.R.F.J., P.A.Z., P.M.P., W.O.S. Visualization: G.U.C., O.R.F.J. Writing – original draft preparation: G.U.C., O.R.F.J., P.A.Z., P.M.P. Writing – review and editing: A.M.D.S., G.U.C., O.R.F.J. Supervision: A.M.D.S. Funding acquisition: A.M.D.S, J.C.S. All authors read, provided critical review, and approved the final manuscript.

## Acknowledgements

We thank Santiago Justo Arevalo and Fernando P.N. Rossi for sharing bioinformatics Python scripts. We also thank Carlos Morais Piroupo for computational support.

