## Supplementary material for "Comparative genomics of *Xylella fastidiosa* suggests determinants of host-specificity and expands its mobile genetic elements repertoire": Table S1; Table S2; Fig S1

**Supplementary Table 1** Features of 132 *X. fastidiosa* RefSeq genome assemblies.

**Supplementary Table 2** List of identified and predicted virulence/pathogenicity factors for *X. fastidiosa*.

**Supplementary Fig. 1** Flowchart describing the workflow implemented to select genomes from the collection of 132 *X. fastidiosa* RefSeq genome assemblies.

| strain name | final collection | assembly accession | genome status | chromossome size (bp) | plasmid number | GC% | number of CDS in chromosome | number of pseudo CDS in chromosome | country of isolation | host of origin | sequence type | completeness % | contamination % |
|---|---|---|---|---|---|---|---|---|---|---|---|---|---|
| 32 | Yes | GCA_000506405 | Contig | 2,607,546 | 0 | 52.45 | 2340 | 133 | Brazil | Coffea-sp | 16 | 99.64 | 0 |
| 3124 | Yes | GCA_001456195 | Complete | 2,748,592 | 0 | 52.63 | 2432 | 148 | Brazil | Coffea-arabica | 16 | 99.64 | 0 |
| 11399 | Yes | GCA_001684415 | Contig | 2,667,741 | 2 | 52.77 | 2452 | 161 | Brazil | Citrus-sinensis | 11 | 99.64 | 0.18 |
| 6c | Yes | GCA_000506905 | Contig | 2,564,403 | 1 | 52.42 | 2358 | 135 | Brazil | Coffea-sp | 14 | 99.28 | 0.18 |
| 9a5c | Yes | GCA_000006725 | Complete | 2,679,305 | 2 | 52.67 | 2455 | 129 | Brazil | Citrus-sinensis | 13 | 99.59 | 0.18 |
| AlmaEM3 | Yes | GCA_018069645 | Complete | 2,487,451 | 0 | 51.76 | 2111 | 104 | USA | Vaccinium-sp | 42 | 99.64 | 0 |
| AlmaEM3 | No | GCA_006369915 | Scaffold | 2,479,954 | 0 | 51.70 | 2136 | 89 | USA | Vaccinium-sp | 42 | 99.64 | 0 |
| Ann-1 | Yes | GCA_000698805 | Complete | 2,750,603 | 1 | 52.10 | 2559 | 158 | USA | Nerium-sp | 5 | 99.64 | 0 |
| Ann-1 | No | GCA_000166855 | Contig | 2,729,755 | 0 | 51.98 | 2624 | 203 | USA | Nerium-sp | 5 | 98.29 | 0.74 |
| ATCC-35871 | Yes | GCA_000428665 | Scaffold | 2,416,255 | 0 | 51.61 | 2092 | 78 | USA | Prunus-sp | 41 | 99.64 | 0 |
| ATCC-35879 | Yes | GCA_011801475 | Complete | 2,565,504 | 1 | 51.82 | 2283 | 127 | USA | Vitis-sp | 2 | 98.91 | 0 |
| ATCC-35879 | No | GCA_000767565 | Contig | 2,522,328 | 0 | 51.79 | 2173 | 85 | USA | Vitis-sp | 2 | 99.23 | 0 |
| B111 | Yes | GCA_013283685 | Scaffold | 2,604,576 | 2 | 52.49 | 2476 | 132 | Brazil | Citrus-sinensis | 11 | 99.64 | 0.18 |
| Bakersfield-1 | Yes | GCA_009664125 | Complete | 2,537,329 | 1 | 51.75 | 2234 | 68 | USA | Vitis-vinifera | 1 | 99.64 | 0 |
| Bakersfield-11 | Yes | GCA_015476015 | Complete | 2,554,506 | 1 | 51.69 | 2254 | 77 | USA | Vitis-vinifera | 1 | 99.64 | 0 |
| Bakersfield-13 | Yes | GCA_015475995 | Complete | 2,537,289 | 1 | 51.75 | 2234 | 76 | USA | Vitis-vinifera | 1 | 99.64 | 0 |
| Bakersfield-14 | Yes | GCA_015475975 | Complete | 2,536,882 | 1 | 51.74 | 2226 | 80 | USA | Vitis-vinifera | 1 | 99.64 | 0 |
| Bakersfield-8 | Yes | GCA_015476035 | Complete | 2,553,893 | 1 | 51.69 | 2262 | 76 | USA | Vitis-vinifera | 1 | 99.64 | 0 |
| BB01 | Yes | GCA_001886315 | Scaffold | 2,511,521 | 0 | 51.77 | 2234 | 99 | USA | Vaccinium-corymbosum | 42 | 99.64 | 0 |
| BB08-1 | Yes | GCA_018069665 | Complete | 2,553,631 | 0 | 51.81 | 2208 | 104 | USA | Vaccinium-sp | 43 | 99.96 | 0 |
| BB08-1 | No | GCA_006369965 | Contig | 2,888,680 | 0 | 51.73 | 2612 | 307 | USA | Vaccinium-sp | 43 | 99.52 | 5.11 |
| BBI64 | No | GCA_006369955 | Contig | 2,927,031 | 0 | 51.79 | 2617 | 292 | USA | Vaccinium-sp | 42 | 98.91 | 7.19 |
| CCPM1 | No | GCA_006370015 | Contig | 2,822,204 | 0 | 51.81 | 2551 | 256 | USA | Vitis-sp | 1 | 99.28 | 3.08 |
| CFBP7969 | No | GCA_004016275 | Contig | 2,431,283 | 0 | 51.50 | 2145 | 69 | USA | Vitis-rotundifolia | 2 | 99.64 | 0 |
| CFBP7970 | Yes | GCA_004016315 | Contig | 2,446,871 | 1 | 51.50 | 2212 | 79 | USA | Vitis-sp | 2 | 99.23 | 0 |
| CFBP8071 | Yes | GCA_004016295 | Contig | 2,446,095 | 1 | 51.53 | 2203 | 80 | USA | Prunus-dulcis | 1 | 99.59 | 0.12 |
| CFBP8072 | Yes | GCA_001469345 | Scaffold | 2,496,662 | 0 | 51.94 | 2310 | 144 | Ecuador | Coffea-arabica | 74 | 99.59 | 0 |
| CFBP8073 | Yes | GCA_001469395 | Scaffold | 2,582,150 | 0 | 51.56 | 2443 | 106 | Mexico | Coffea-canephora | 75 | 99.63 | 0 |
| CFBP8078 | Yes | GCA_004016365 | Contig | 2,596,546 | 0 | 51.69 | 2414 | 128 | USA | Vinca-sp | 51 | 99.64 | 0 |
| CFBP8082 | Yes | GCA_004016375 | Contig | 2,525,300 | 1 | 51.53 | 2286 | 79 | USA | Ambrosia-artenisiifolia | 2 | 99.64 | 0 |
| CFBP8351 | Yes | GCA_004016405 | Contig | 2,440,242 | 1 | 51.50 | 2199 | 81 | USA | Vitis-vinifera | 1 | 99.64 | 0 |
| CFBP8356 | Yes | GCA_004016415 | Scaffold | 2,536,149 | 0 | 51.59 | 2308 | 114 | Costa-Rica | Coffea-arabica | 76 | 99.28 | 0 |
| CFBP8416 | Yes | GCA_001971475 | Contig | 2,466,748 | 0 | 51.79 | 2212 | 344 | France | Polygala-myrtifolia | 7 | 98.31 | 0.02 |
| CFBP8417 | Yes | GCA_001971505 | Contig | 2,504,981 | 0 | 51.85 | 2325 | 76 | France | Spartium-junceum | 6 | 99.64 | 0 |
| CFBP8418 | No | GCA_001971465 | Contig | 2,513,969 | 0 | 51.88 | 2357 | 78 | France | Spartium-junceum | 6 | 99.64 | 0 |
| CO33 | Yes | GCA_001417925 | Contig | 2,681,926 | 0 | 51.70 | 2415 | 122 | Costa-Rica | Coffea-sp | 72 | 99.28 | 0 |
| CoDiRO | Yes | GCA_000811965 | Contig | 2,507,614 | 1 | 52.03 | 2215 | 123 | Italy | Olea-sp | 53 | 99.64 | 0 |
| COF0324 | Yes | GCA_001549815 | Contig | 2,702,997 | 4 | 52.45 | 2619 | 123 | Brazil | Coffea-sp | 14 | 99.64 | 0.18 |
| COF0407 | Yes | GCA_001549825 | Contig | 2,463,947 | 4 | 51.93 | 2312 | 119 | Costa-Rica | Coffea-sp | 53 | 99.28 | 0 |
| CVC0251 | Yes | GCA_001549765 | Contig | 2,625,257 | 4 | 52.57 | 2558 | 129 | Brazil | Citrus-sinensis | 11 | 99.64 | 0.18 |
| CVC0256 | Yes | GCA_001549745 | Contig | 2,587,189 | 4 | 52.60 | 2533 | 126 | Brazil | Citrus-sinensis | 11 | 98.91 | 0.18 |
| De-Donno | Yes | GCA_002117875 | Complete | 2,508,465 | 1 | 51.99 | 2223 | 121 | Italy | Olea-europaea | 53 | 99.59 | 0 |
| Dixon | Yes | GCA_000166835 | Scaffold | 2,582,607 | 1 | 52.07 | 2366 | 96 | USA | Prunus-sp | 6 | 99.63 | 0 |
| DSM-10026 | Yes | GCA_900129695 | Scaffold | 2,431,652 | 0 | 51.35 | 2115 | 71 | USA | Vitis-vinifera | 2 | 99.23 | 0 |
| EB92-1 | Yes | GCA_000219235 | Contig | 2,434,000 | 1 | 51.52 | 2236 | 71 | USA | Sambucus-canadensis | 1 | 99.64 | 0.18 |
| ESVL | Yes | GCA_004023385 | Contig | 2,493,528 | 2 | 51.77 | 2296 | 98 | Spain | Prunus-dulcis | 6 | 99.64 | 0.36 |
| Fb7 | Yes | GCA_001456335 | Complete | 2,659,912 | 1 | 52.54 | 2447 | 273 | Argentina | Citrus-sp | 69 | 98.78 | 0 |
| Fillmore | Yes | GCA_012974105 | Complete | 2,526,544 | 0 | 51.98 | 2195 | 98 | USA | Olea-europaea | 81 | 99.64 | 0 |
| GB514 | Yes | GCA_000148405 | Complete | 2,491,203 | 1 | 51.78 | 2169 | 194 | USA | Vitis-sp | 1 | 99.18 | 0 |
| Griffin-1 | Yes | GCA_000466025 | Contig | 2,387,314 | 0 | 51.73 | 2088 | 193 | USA | Quercus-rubra | 7 | 98.46 | 0 |
| GV156 | Yes | GCA_009910885 | Contig | 2,521,515 | 0 | 51.74 | 2190 | 106 | Taiwan | Vitis-vinifera | 2 | 99.46 | 0 |
| GV230 | Yes | GCA_014249995 | Complete | 2,514,993 | 0 | 51.74 | 2159 | 63 | Taiwan | Vitis-labrusca | 2 | 99.64 | 0 |
| Hib4 | Yes | GCA_001456315 | Complete | 2,813,286 | 1 | 52.70 | 2594 | 142 | Brazil | Hibiscus-sp | 70 | 99.64 | 1.45 |
| IAS-AXF212H7 | Yes | GCA_009669445 | Contig | 2,521,176 | 1 | 51.75 | 2359 | 99 | Spain | Prunus-dulcis | 6 | 99.64 | 0.41 |
| IAS-AXF-235T10 | Yes | GCA_009669465 | Contig | 2,548,843 | 1 | 51.74 | 2424 | 115 | Spain | Prunus-dulcis | 6 | 99.64 | 0.11 |
| IVIA5235 | Yes | GCA_003515915 | Complete | 2,537,917 | 1 | 51.73 | 2224 | 66 | Spain | Prunus-avium | 1 | 99.64 | 0 |
| IVIA5901 | Yes | GCA_004023395 | Complete | 2,559,157 | 0 | 51.98 | 2231 | 123 | Spain | Prunus-dulcis | 6 | 99.64 | 0 |
| IVIA6586-2 | Yes | GCA_009669335 | Contig | 2,518,561 | 2 | 51.76 | 2397 | 96 | Spain | Helicrysum-italicum | 6 | 99.64 | 0.72 |
| IVIA6629 | No | GCA_009669365 | Contig | 2,730,337 | 1 | 53.17 | 2941 | 100 | Spain | Rhamnus-alaternus | 6 | 99.64 | 1.16 |
| IVIA6731 | Yes | GCA_009669375 | Contig | 2,539,193 | 2 | 51.91 | 2474 | 99 | Spain | Helicrysum-italicum | 6 | 99.64 | 1.13 |
| IVIA6902 | No | GCA_009669405 | Contig | 2,870,108 | 0 | 53.48 | 3337 | 141 | Spain | Prunus-dulcis | 6 | 99.64 | 2.05 |
| IVIA6903 | No | GCA_009669425 | Contig | 2,602,829 | 0 | 52.12 | 2556 | 104 | Spain | Prunus-dulcis | 6 | 99.64 | 0.97 |

| | | | | | | | | | | | | | |
|---|---|---|---|---|---|---|---|---|---|---|---|---|---|
| J1a12 | Yes | GCA_001456235 | Complete | 2,788,789 | 2 | 52.91 | 2641 | 207 | Brazil | Citrus-sp | 11 | 99.59 | 0.18 |
| KLN59-3-GFP-Temecula1 | No | GCA_006369995 | Contig | 2,459,746 | 0 | 51.57 | 2200 | 62 | USA | Lab-variant | 1 | 99.64 | 0 |
| LM10 | Yes | GCA_012974145 | Complete | 2,669,650 | 0 | 52.12 | 2375 | 113 | USA | Olea-europaea | 7 | 99.64 | 0 |
| M12 | Yes | GCA_000019325 | Complete | 2,475,130 | 0 | 51.92 | 2123 | 93 | USA | Prunus-sp | 7 | 99.64 | 0 |
| M23 | Yes | GCA_000019765 | Complete | 2,535,690 | 1 | 51.76 | 2228 | 69 | USA | Prunus-sp | 1 | 99.64 | 0 |
| Ma1 | No | GCA_018449155 | Contig | 2,414,358 | 0 | 51.70 | 2101 | 91 | Italy | Rhamnus-alaternus | 87 | 99.64 | 0 |
| Ma10 | No | GCA_018449235 | Contig | 2,415,263 | 0 | 51.70 | 2098 | 91 | Italy | Rhamnus-alaternus | 87 | 99.64 | 0 |
| Ma151 | Yes | GCA_018449095 | Contig | 2,416,213 | 0 | 51.69 | 2111 | 89 | Italy | Rhamnus-alaternus | 87 | 99.64 | 0 |
| Ma185 | No | GCA_018449105 | Contig | 2,412,610 | 0 | 51.69 | 2093 | 87 | Italy | Polygala-myrtifolia | 87 | 99.64 | 0 |
| Ma26 | No | GCA_018449175 | Contig | 2,412,928 | 0 | 51.70 | 2094 | 89 | Italy | Spartium-junceum | 87 | 99.64 | 0 |
| Ma29 | No | GCA_018449135 | Contig | 2,426,899 | 0 | 51.69 | 2100 | 91 | Italy | Prunus-dulcis | 87 | 99.64 | 0.72 |
| MUL0034 | Yes | GCA_000698825 | Complete | 2,642,186 | 1 | 51.97 | 2388 | 129 | USA | Morus-alba | 30 | 99.64 | 0 |
| Mul-MD | Yes | GCA_000567985 | Contig | 2,520,555 | 0 | 51.65 | 2272 | 119 | USA | Morus-alba | 29 | 99.64 | 0 |
| Mus-1 | No | GCA_009812445 | Contig | 2,438,200 | 0 | 51.51 | 2156 | 223 | USA | Vitis-sp | 2 | 99.29 | 0.36 |
| NOB1 | Yes | GCA_012952075 | Scaffold | 2,418,556 | 0 | 51.46 | 2123 | 68 | USA | Vitis-rotundifolia | 2 | 99.64 | 0 |
| NS1-CmR-TemeculaL | No | GCA_006370045 | Scaffold | 2,510,201 | 0 | 51.44 | 2338 | 86 | USA | Lab-variant | 1 | 99.64 | 1.45 |
| OK3 | Yes | GCA_012952085 | Scaffold | 2,415,390 | 0 | 51.48 | 2119 | 68 | USA | Vitis-vinifera | 2 | 99.64 | 0 |
| OLS0478 | Yes | GCA_001549755 | Contig | 2,483,278 | 2 | 51.95 | 2291 | 120 | Costa-Rica | Nerium-sp | 53 | 99.64 | 0 |
| OLS0479 | Yes | GCA_001549735 | Contig | 2,474,529 | 4 | 51.96 | 2338 | 116 | Costa-Rica | Nerium-sp | 53 | 99.23 | 0.36 |
| PD7202 | Yes | GCA_006370235 | Contig | 2,629,977 | 0 | 52.10 | 2354 | 141 | Netherlands | Plant-tissue | undetermined | 98.14 | 0 |
| PD7211 | Yes | GCA_006370175 | Contig | 2,651,656 | 0 | 52.16 | 2336 | 137 | Netherlands | Plant-tissue | 73 | 99.64 | 0 |
| pglA-KmR-Fetzer | No | GCA_006370115 | Scaffold | 2,467,450 | 1 | 51.51 | 2246 | 89 | USA | Lab-variant | 4 | 99.64 | 0 |
| Pr8x | Yes | GCA_001456295 | Complete | 2,666,240 | 1 | 52.63 | 2423 | 134 | Brazil | Prunus-sp | 14 | 99.59 | 0.18 |
| RAAR14-plum327 | Yes | GCA_009695495 | Contig | 2,543,559 | 0 | 51.64 | 2340 | 120 | Brazil | Prunus-domestica | 26 | 99.64 | 0.18 |
| RAAR6-Butte | Yes | GCA_009695485 | Contig | 2,466,226 | 0 | 51.81 | 2196 | 76 | USA | Prunus-dulcis | 7 | 99.64 | 0 |
| Red-Oak-2 | Yes | GCA_015475935 | Complete | 2,476,446 | 0 | 51.92 | 2117 | 96 | USA | Quercus-rubra | 7 | 99.64 | 0 |
| RH1 | Yes | GCA_012974125 | Complete | 2,678,425 | 0 | 52.15 | 2388 | 123 | USA | Olea-europaea | 7 | 99.64 | 1.45 |
| Riv5 | Yes | GCA_015475955 | Complete | 2,499,994 | 1 | 51.96 | 2175 | 94 | USA | Prunus-cerasifera | 34 | 99.64 | 0 |
| Salento-1 | Yes | GCA_002954185 | Complete | 2,508,097 | 1 | 51.98 | 2231 | 230 | Italy | Olea-europaea | 53 | 99.11 | 0 |
| Salento-2 | Yes | GCA_002954205 | Complete | 2,508,296 | 1 | 51.98 | 2232 | 182 | Italy | Olea-europaea | 53 | 99.59 | 0 |
| Stag-s-Leap | Yes | GCA_001572105 | Contig | 2,489,133 | 1 | 51.73 | 2161 | 80 | USA | Vitis-sp | 1 | 99.64 | 0 |
| sycamore-Sy-VA | Yes | GCA_000732705 | Contig | 2,475,880 | 0 | 51.64 | 2211 | 118 | USA | Platanus-occidentalis | 8 | 99.64 | 0.09 |
| Temecula1 | Yes | GCA_000007245 | Complete | 2,519,802 | 1 | 51.78 | 2172 | 64 | USA | Vitis-sp | 1 | 99.64 | 0 |
| Temecula1Star | Yes | GCA_006370185 | Contig | 2,454,666 | 0 | 51.57 | 2190 | 62 | USA | Vitis-sp | 1 | 99.64 | 0 |
| TemeculaL | Yes | GCA_006370155 | Scaffold | 2,536,542 | 0 | 51.46 | 2290 | 86 | USA | Vitis-sp | 1 | 99.64 | 0 |
| TemL-Alma-Rec1 | No | GCA_006369895 | Scaffold | 2,522,659 | 0 | 51.47 | 2262 | 79 | USA | Lab-variant | 1 | 99.64 | 0 |
| TemL-Alma-Rec2 | No | GCA_006369905 | Scaffold | 2,654,426 | 0 | 51.02 | 2396 | 81 | USA | Lab-variant | 1 | 99.64 | 0.04 |
| TemL-NS1pglA-Rec1 | No | GCA_006370085 | Scaffold | 2,525,255 | 0 | 51.47 | 2278 | 88 | USA | Lab-variant | 1 | 99.64 | 0 |
| TOS14 | Yes | GCA_007713995 | Contig | 2,445,518 | 0 | 51.74 | 2140 | 96 | Italy | Spartium-junceum | 87 | 99.64 | 0 |
| TOS4 | Yes | GCA_007713905 | Contig | 2,445,114 | 0 | 51.73 | 2139 | 95 | Italy | Prunus-dulcis | 87 | 99.64 | 0.36 |
| TOS5 | Yes | GCA_007713945 | Contig | 2,443,867 | 0 | 51.73 | 2131 | 94 | Italy | Polygala-myrtifolia | 87 | 99.64 | 0 |
| TPD3 | Yes | GCA_007845655 | Contig | 2,422,083 | 0 | 51.49 | 2241 | 58 | Taiwan | Vitis-vinifera | 2 | 99.64 | 0.36 |
| TPD4 | Yes | GCA_007845705 | Contig | 2,427,175 | 0 | 51.47 | 2256 | 59 | Taiwan | Vitis-vinifera | 2 | 99.64 | 0.21 |
| U24D | Yes | GCA_001456275 | Complete | 2,681,334 | 1 | 52.68 | 2451 | 160 | Brazil | Citrus-sinensis | 13 | 99.59 | 0.18 |
| VB11 | Yes | GCA_012952095 | Scaffold | 2,423,256 | 1 | 51.43 | 2166 | 81 | USA | Vitis-vinifera | 2 | 99.64 | 0.01 |
| WM1-1 | Yes | GCA_006370215 | Contig | 2,470,027 | 0 | 51.59 | 2151 | 67 | USA | Vitis-sp | 2 | 99.64 | 0 |
| WM1-1-GFP-Rec1 | No | GCA_006370035 | Scaffold | 2,448,134 | 0 | 51.49 | 2186 | 62 | USA | Lab-variant | 2 | 99.64 | 0 |
| WM1-1-GFP-Rec2 | No | GCA_006370055 | Scaffold | 2,452,477 | 0 | 51.49 | 2200 | 67 | USA | Lab-variant | 2 | 99.64 | 0 |
| XF3348 | Yes | GCA_009669515 | Contig | 2,572,158 | 1 | 51.81 | 2459 | 105 | Spain | Prunus-dulcis | 81 | 99.64 | 1.37 |
| XF-3960-18 | No | GCA_014856905 | Scaffold | 2,518,024 | 0 | 51.79 | 2287 | 89 | Spain | Prunus-dulcis | 81 | 99.64 | 0 |
| XF-3967-18 | No | GCA_015102005 | Scaffold | 2,484,418 | 0 | 51.51 | 2230 | 68 | Spain | Vitis-vinifera | 1 | 99.64 | 0.36 |
| XRB | Yes | GCA_013283695 | Scaffold | 2,628,251 | 2 | 52.53 | 2453 | 128 | Brazil | Citrus-sp | 11 | 99.59 | 0.18 |
| XYL1732 | Yes | GCA_003973705 | Contig | 2,444,109 | 0 | 51.45 | 2185 | 62 | Spain | Vitis-sp | 1 | 99.64 | 0 |
| XYL1752 | Yes | GCA_009669505 | Contig | 2,578,009 | 1 | 51.82 | 2469 | 99 | Spain | Prunus-dulcis | 81 | 99.64 | 0.25 |
| XYL1966-18 | No | GCA_014856865 | Scaffold | 2,514,978 | 0 | 51.79 | 2287 | 88 | Spain | Olea-europaea | 81 | 99.64 | 0 |
| XYL1968-18 | Yes | GCA_014856935 | Contig | 2,518,610 | 0 | 51.80 | 2288 | 81 | Spain | Olea-europaea | 81 | 99.64 | 0 |
| XYL1978-18 | No | GCA_014856645 | Scaffold | 2,484,291 | 0 | 51.50 | 2235 | 67 | Spain | Prunus-dulcis | 1 | 99.64 | 0 |
| XYL1980-18 | No | GCA_014856695 | Scaffold | 2,485,332 | 0 | 51.51 | 2221 | 61 | Spain | Prunus-dulcis | 1 | 99.64 | 0 |
| XYL1981 | Yes | GCA_009669455 | Contig | 2,554,510 | 0 | 51.83 | 2414 | 99 | Spain | Ficus-carica | 81 | 99.64 | 0.12 |
| XYL1981-18 | No | GCA_014856675 | Scaffold | 2,509,949 | 0 | 51.80 | 2279 | 89 | Spain | Prunus-dulcis | 81 | 99.64 | 0 |
| XYL2014-18 | No | GCA_014856685 | Scaffold | 2,484,228 | 0 | 51.50 | 2225 | 64 | Spain | Prunus-dulcis | 1 | 99.64 | 0 |
| XYL2017-18 | No | GCA_014856735 | Scaffold | 2,482,742 | 0 | 51.49 | 2219 | 66 | Spain | Prunus-dulcis | 1 | 99.64 | 0 |
| XYL2055 | Yes | GCA_003973695 | Contig | 2,456,780 | 0 | 51.46 | 2197 | 67 | Spain | Vitis-sp | 1 | 99.64 | 0.91 |
| XYL2093-18 | No | GCA_014856745 | Scaffold | 2,486,223 | 0 | 51.51 | 2225 | 61 | Spain | Prunus-dulcis | 1 | 99.64 | 0.36 |

| | | | | | | | | | | | | | |
|---|---|---|---|---|---|---|---|---|---|---|---|---|---|
| XYL2106-18 | No | GCA_014856775 | Scaffold | 2,484,940 | 0 | 51.50 | 2224 | 65 | Spain | Prunus-dulcis | 1 | 99.64 | 0.36 |
| XYL2107-18 | Yes | GCA_014856795 | Scaffold | 2,483,447 | 0 | 51.50 | 2222 | 63 | Spain | Prunus-dulcis | 1 | 99.64 | 0.36 |
| XYL2153-18 | Yes | GCA_014856785 | Scaffold | 2,484,994 | 0 | 51.51 | 2222 | 65 | Spain | Vitis-vinifera | 1 | 99.64 | 0 |
| XYL2177-18 | No | GCA_014856835 | Scaffold | 2,484,593 | 0 | 51.50 | 2230 | 60 | Spain | Vitis-vinifera | 1 | 99.64 | 0 |
| XYL2400-18 | No | GCA_014856845 | Scaffold | 2,486,358 | 0 | 51.50 | 2236 | 64 | Spain | Vitis-vinifera | 1 | 99.64 | 0 |
| XYL2508-18 | No | GCA_014856895 | Scaffold | 2,485,230 | 0 | 51.51 | 2224 | 63 | Spain | Vitis-vinifera | 1 | 99.64 | 0.36 |

| Gene ID in Temecula1 strain | Gene | Description | Function | Number of genomes with intact gene |
|---|---|---|---|---|
| PD0023 | | PilY1 homolog | Twitching motility | 94 |
| PD0058 | *fimF* | Fimbrial adhesin: cell-cell aggregation, biofilm formation | Adhesin | 94 |
| PD0062 | *fimA* | Fimbrial adhesin: cell-cell aggregation, biofilm formation | Adhesin | 94 |
| PD0218 | | Serine protease autotransporter | Protease | 94 |
| PD0233 | *rpfB* | DSF synthesis, quorum sensing | Quorum sensing | 86 |
| PD0279 | *cgsA* | Cyclic di-GMP synthase A, quorum sensing | Quorum sensing | 90 |
| PD0313 | | serine protease/autotransporter-associated beta strand repeat protein | Hydrolytic enzyme | 94 |
| PD0406 | *rpfC* | Response regulator, quorum sensing | Quorum sensing | 92 |
| PD0407 | *rpfF* | DSF synthase, quorum sensing | Quorum sensing | 94 |
| PD0502 | | PilY1 homolog | Twitching motility | 94 |
| PD0519 | | Acetyl esterase/lipase | Hydrolytic enzyme | 89 |
| PD0528 | *xatA* | Autotransporter beta- domain protein | Adhesin | 79 |
| PD0529 | *cbsA* | 1,4-beta-cellobiosidase | Hydrolytic enzyme | 82 |
| PD0639 | *LptA* | Lipopolysaccharide transport periplasmic protein LptA | Adhesin | 94 |
| PD0657 | | Putative extracellular serine protease | Protease | 80 |
| PD0731 | *xadA1* | Afimbrial adhesin: cell surface attachment, biofilm formation | Adhesin | 92 |
| PD0732 | *xpsE* | Type II secretion system: ATPase | Secretion system | 94 |
| PD0744 | *xadA2* | Afimbrial adhesin | Adhesin | 88 |
| PD0794 | *xatC* | Uncharacterized membrane protein | Membrane protein | 94 |
| PD0814 | *wzy* | O-antigen ligase family protein | Polysaccharide synthesis | 93 |
| PD0815 | | Glycosyltransferase family 2 protein | Polysaccharide synthesis | 93 |
| PD0816 | | CDP-glycerol glycerophosphotransferase family protein | Polysaccharide synthesis | 94 |
| PD0824 | *xadA3* | Afimbrial adhesin | Adhesin | 88 |
| PD0843 | *tonB1* | Iron and vitamin B12 transport | Membrane protein | 94 |
| PD0848 | *pilL* | Type IV pilus: regulates twitching | Regulatory system | 93 |
| PD0855 | | VirK protein | Pathogenesis | 94 |
| PD0915 | *zot* | Putative Zot-like toxin - filamentous phage Cf1c related protein | Toxin | 55 |
| PD0928 | *zot* | Putative Zot-like toxin - filamentous phage Cf1c related protein | Toxin | 55 |
| PD0950 | | Serine protease/autotransporter-associated beta strand repeat protein | Hydrolytic enzyme | 94 |
| PD0956 | *prtA* | Serine protease | Protease | 52 |
| PD0985 | | DUF769 domain-containing protein | Hypothetical protein | 94 |
| PD0986 | | Putative haemagglutinin-like protein | Adhesin | 72 |
| PD1099 | *dinJ/relE* | Toxin-anti-toxin system, regulatory system | Regulatory system | 70 |
| PD1100 | | mRNA-degrading endonuclease (mRNA interferase) YafQ toxin of type II toxin-anti | Regulatory system | 70 |
| PD1211 | | Putative secreted lipase (lipases/esterases) | Hydrolytic enzyme | 94 |
| PD1249 | | Hemagglutinin-like secreted protein | Adhesin | 94 |
| PD1284 | *algU* | Alternate sigma factor: pathogenicity-related genes, stress response | Regulatory system | 94 |
| PD1311 | | Acyl-coenzyme A (acyl-CoA) synthetase | Metabolism | 93 |
| PD1379 | *xatB* | Autotransporter beta- domain protein | Adhesin | 94 |
| PD1380 | *csp1* | Cold shock protein | Stress response | 94 |
| PD1386 | *xhpT* | Two-component regulatory system: pathogenicity-related genes, EPS synthesis | Regulatory system | 93 |

| | | | | |
|---|---|---|---|---|
| PD1391 | *gumH* | Exopolysaccharide synthesis | Polysaccharide synthesis | 92 |
| PD1394 | *gumD* | Exopolysaccharide synthesis | Polysaccharide synthesis | 91 |
| PD1403 | | Multidrug efflux pump | Membrane protein | 94 |
| PD1427 | | Calcium binding protein - RTX toxin-related | Bacterial toxicity | 94 |
| PD1452 | | Lipopolysaccharide core biosynthesis protein | Polysaccharide synthesis | 92 |
| PD1485 | *pglA* | Polygalacturonase: pit membrane degradation | Hydrolytic enzyme | 82 |
| PD1506 | | Hemolysin-type calcium binding domain protein | Pathogenesis | 94 |
| PD1611 | *pilY* | type IV pilus assembly protein PilY1 | Twitching motility | 91 |
| PD1640 | | Beta-glucosidase | Hydrolytic enzyme | 94 |
| PD1671 | | Two-component regulatory system: pathogenicity-related genes | Regulatory system | 94 |
| PD1678 | *phoQ* | Two-component regulatory system: pathogenicity-related genes, survival genes | Regulatory system | 94 |
| PD1679 | *phoP* | Two-component regulatory system: pathogenicity-related genes, survival genes | Regulatory system | 94 |
| PD1702 | *lesB* | Lipase/esterase | Hydrolytic enzyme | 94 |
| PD1703 | *lesA* | Lipase/esterase | Hydrolytic enzyme | 94 |
| PD1792 | *hxfB* | Haemagglutinin: cell-cell aggregation | Adhesin | 94 |
| PD1801 | | Lipopolysaccharide biosynthesis protein (transferase) | Polysaccharide synthesis | 94 |
| PD1826 | *chiA* | Chitinase | Hydrolytic enzyme | 93 |
| PD1833 | | beta-galactosidase | Hydrolytic enzyme | 94 |
| PD1850 | | Peptidase (M20/M25/M40 family) | Hydrolytic enzyme | 94 |
| PD1851 | *engXCA2* | Endoglucanase: pit membrane degradation | Hydrolytic enzyme | 93 |
| PD1856 | *engXCA1* | Endoglucanase: pit membrane degradation | Hydrolytic enzyme | 93 |
| PD1879 | | Lipase/esterase | Hydrolytic enzyme | 94 |
| PD1924 | *pilA1* | Pilin protein - long type IV pili | Twitching motility | 94 |
| PD1926 | *pilA2* | Pilin protein - long type IV pili | Twitching motility | 94 |
| PD1964 | *tolC* | Type I secretion system | Secretion system | 94 |
| PD1984 | *gacA* | Two-component regulatory system: pathogenicity-related genes | Regulatory system | 92 |
| PD2061 | | endo-1,4-beta-glucanase | Hydrolytic enzyme | 94 |
| PD2084 | | DUF769 domain-containing protein | Hypothetical protein | 65 |
| PD2118 | *hxfA* | Haemagglutinin: cell-cell aggregation | Adhesin | 94 |

### Genome selection

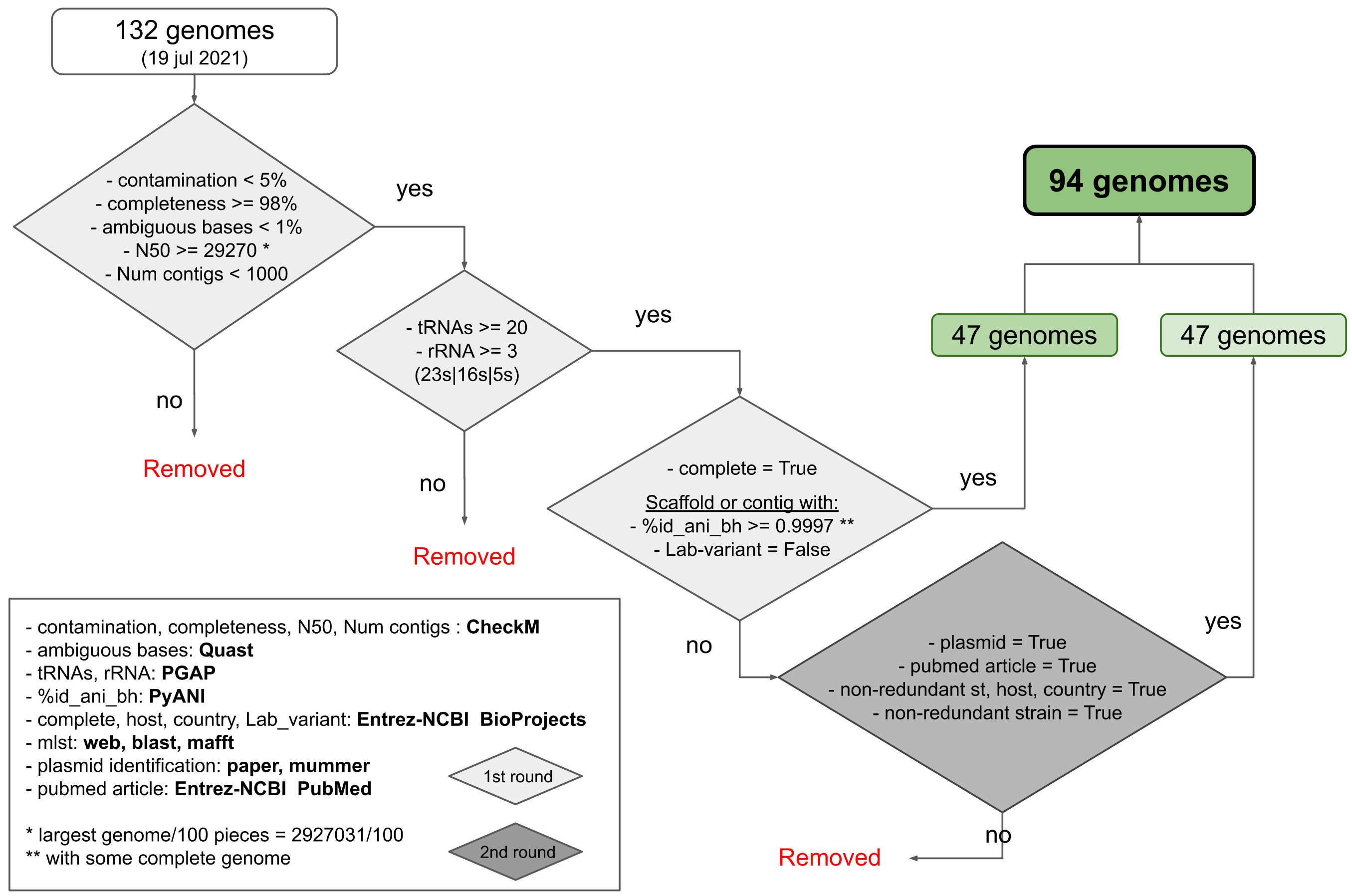


Fig. S1
